# From Randomness to Recognition: Modeling the Evolution of DNA Sequence Information During SELEX

**DOI:** 10.1101/2025.03.31.646277

**Authors:** Varun Maher, Daniel Martin, David Spetzler, Zhan-Gong Zhao, Heather O’Neill, Neal W. Woodbury

## Abstract

<u>S</u>ystematic <u>E</u>volution of <u>L</u>igands through <u>E</u>xponential <u>E</u>nrichment (SELEX) was used as a model system to explore the evolution of DNA sequence information and function during enrichment of molecular recognition to a series of related target molecules. Using a Natural Language Processing (NLP) based approach, a model was trained on an unlabeled mixture of oligonucleotide sequences sampled from both unenriched libraries and from six libraries that had been enriched for binding to either peptide or protein targets. This general model (pre-trained model) was then used to generate latent space representations of sequences that contained embedded binding information. Unsupervised clustering was used to create two dimensional maps that allowed comparison of the overlap between the latent space representations of sequences from different unenriched and enriched libraries. Replicate, independent enrichments to the same targets starting from completely unique random libraries gave rise to essentially indistinguishable latent space representations. However, similar representations between unenriched and enriched sequences, or between enriched sequences from different targets, resulted in distinct clustering patterns. The extent of overlap between the patterns from unenriched and enriched libraries was correlated with overall target binding by the enriched library. Further, the pre-trained model was fine-tuned to classify which target library each sequence belonged to, and the accuracy of classification between unenriched and enriched sequences was also correlated with the overall binding of the enriched library to its target. Finally, it was possible to distinguish libraries enriched for binding to two different targets differing by as few as 1 in 30 amino acids. Thus, using NLP-based approaches, it is possible to relate the DNA sequences present in an enriched library with the ability of the library to specifically bind its intended target, providing a tool for both guiding the enrichment process and more deeply understanding the structure-binding relationships that evolve.

## Author Summary

The ability to select DNA sequences that recognize and bind specific molecules has been applied to many biological and medical applications, from drug development to diagnostics. In this study, the evolution of DNA sequences during the process of selecting high-affinity binders to target molecules was explored. A machine learning model was trained on a mixture of sequences from libraries that had been enriched for binding to different targets, as well as from unenriched libraries. This model captured hidden patterns in the sequences, making it possible to create maps that visually represent how sequences change as they evolve to recognize their targets. Remarkably, independent experiments that started with completely unique sequences produced nearly identical patterns when enriched for the same target, demonstrating the reproducibility of molecular evolution from a functional perspective. Additionally, it was possible to distinguish between libraries enriched for closely related targets, even when they differed by only a single amino acid. These findings show that machine learning models can provide insights into the molecular evolution of DNA sequences and can serve both as tools for optimizing their binding properties and providing a deeper understanding of sequence characteristics that are important for binding to specific targets.

## Introduction

The development of aptamers or enriched oligodeoxynucleotide sequences through Systematic Evolution of Ligands by Exponential Enrichment (SELEX) involves iterative cycles where a large, diverse library of sequences (10^8^ – 10^15^ unique species) is progressively enriched for sequences that bind specifically to a target of interest. Since its introduction^1,2^, SELEX has been used to generate aptamers for diverse targets, including organic dyes^2^, small molecules^3–5^, proteins^6–10^, tumor marker peptides^11^, and complex targets such as cells or lysates from FFPE tissues^12–14^. Its simplicity makes it an excellent model system for exploring how chemical information changes when starting a process with random structures and evolving towards specific activities like ligand binding. Recent advances in applying machine learning and natural language processing methods to macromolecules like proteins and nucleic acids have made it possible to create meaningful representations of sequence that incorporate functional capability. This in turn opens the door for developing new tools for measuring and characterizing the information content of molecules and functionally enriched molecular libraries. The SELEX process itself could benefit from a means of monitoring functional information content during enrichment, allowing one to systematically address challenges such as the retention of non-specific sequences, incomplete washing after binding, preferential amplification of suboptimal binders to the target^15,16^, and an inability to fully explore the entire combinatorial space of possible oligonucleotides. Hence, enriched libraries can be quite heterogeneous, containing both binding and nonbinding sequences, which reduces binding specificity of the library to the target and complicates identification of specific aptamers. There would be both practical and intellectual value to having a means of estimating target specific information content of sequences identified by Next-Generation Sequencing (NGS) from enriched libraries during the SELEX process by modeling the profile of sequences detected at any point during the enrichment process. Monitoring the level of target specific information during enrichment may enable faster protocol optimization, requiring fewer rounds of enrichment, particularly if information content early in the selection is indicative of the outcome, and potentially helping reduced downstream validation of irrelevant sequences. The ability to model the quantitative relationship between targets and enriched sequence information content could lead to a deeper understanding of how target structure drives sequence evolution during the enrichment process and provide a paradigm for the evolution of chemical information for generally using a well-characterized model system.

Various computational approaches have been used to identify functional aptamer sequences, either directly *in silico* or from high-throughput SELEX data obtained during the enrichment process, with notable success in identifying and optimizing aptamer sequences. This has been the subject of several recent reviews^17–19^. Many of these approaches are predicated on identifying potential aptamer sequences by clustering enriched sequences based on similarity, searching for motifs and common n-mers, and scoring candidates on criteria such as binding affinity and secondary structure stability^20–24^. The use of machine learning and deep learning models, such as supervised learning classifiers and convolutional neural networks to predict potential aptamer binding based on sequence features, effectively allows screening of large datasets to identify potential high-affinity candidates^25,26^. Other examples include SMART-Aptamer^27^ and RaptRanker^23^, which rank aptamers based on sequence abundance, motif stability, and predicted structure.

In addition to identification, computational methods are increasingly used for potential aptamer sequence optimization. A few tools have been developed for secondary (2D) and tertiary (3D) structure prediction, such as Mfold^28^ and RNAComposer^29^, and these provide metrics that can be used in optimizing aptamer-target interactions at a molecular level. In addition, a few researchers are integrating molecular docking and molecular dynamics simulations into aptamer optimization protocols^30,31^. Tools such as AutoDock^32^ and GROMACS^33^ assess the stability and binding energies of aptamer-target complexes, and this facilitates further optimization through iterative sequence modifications.

The approaches described above mostly focus on finding discrete motifs or other sequence patterns, often augmented by either structural predictions or binding predictions. The success of large language models, and their counterparts in the analysis of biological polymers^34,35^, suggests that it should be possible to look at the sequence space of enriched libraries in a similar fashion, developing latent space representations of sequences related to enrichment towards binding to a particular target. Natural Language Processing (NLP) has shown promise in building biological sequence-based embeddings that can be used as latent space representations to integrate information for use in mapping the biological properties of the sequences themselves.Modern NLP works by building representations of words or sequences that capture their meaning relative to other words. This type of representation learning allows the exploration of features of raw data in an unlabeled fashion, uncovering complex associations that are otherwise difficult to identify. Biological sequences vectorized by representation learning have successfully been used in tasks such as structure prediction^35,36^ and function association^37^ using similarity and differences within these sequences in representation space. Deep learning approaches with raw protein sequences have shown promise in creating models and embeddings through representation learning which are then used to extract features for predictive tasks such as cell compartmentalization of proteins and predicting biological properties of engineered sequences^38^. Additionally, learned representations offer an opportunity to visualize groups of sequences with similar features in latent space after dimension reduction. In previous sequence library analysis using dimension reduction, a variational autoencoder was used to create an embedding for in-silico aptamer generation where simulated sequence data was found to be distributed in latent space based on motif information^39^. Sequence embeddings were created using two selection datasets against separate targets and this was then used to successfully generate truncated sequences also capable of binding the targets. More recently, the use of latent space representation has been extended to the development of a diffusion model for generative prediction^40^.

Here, latent space sequence representations based on SELEX data are created using a masked language model (MLM) with bidirectional Long Short-Term Memory (biLSTM) architecture, a type of recurrent neural network. This approach is similar to one that has been used previously to successfully generate protein sequence embeddings^41^. These latent space sequence representations are then interrogated for their ability to effectively capture information that is correlated with biological properties of the sequences, such as binding affinity to the enrichment target and specificity for one target vs another in targets with varying degrees of similarity. A key aspect of this approach is its ability to obtain target-specific information simply by characterizing the enriched sequences, without explicitly incorporating binding or target information during the self-supervised training of the model. Using this approach , several fundamental questions can be explored. Do two independent enrichments using the same target result in equivalent latent space representations? Are the latent space representations specific enough to differentiate one target from another? How similar can two targets be and still create different latent space representations? Is there enough target-specific information in the latent space representation to predict binding to the target? The goal of this study is to lay a foundation for using these models both as a predictive tool that can guide the SELEX process by measuring target-specific information content during the enrichment process and as a means of exploring the molecular recognition relationship between target structure and sequence content in enriched libraries.

## Results

A schematic of the overall analysis workflow is depicted in **Fig 1**. Random single-stranded oligodeoxynucleotide libraries (approximately 4.0 x 10^12 molecules/target initially) were used to enrich on any of six protein or peptide targets by performing six rounds of binding, PCR amplification, and target binding ssDNA strand isolation in a high throughput workflow using targets conjugated to magnetic beads. (**Fig 1A**). The starting unenriched libraries and final enriched libraries (approximately 35 x 10^9 molecules per sample in a normalized pool) were flowed over a flowcell for sequencing by NGS. A random sampling of unlabeled sequences from all unenriched and enriched libraries (approximately 12% of the sequenced space) were used as the inputs to a masked language model resulting in latent sequence representations and a pre-trained overall model (**Fig 1B**). Those latent sequence representations were used in unsupervised clustering analysis and the pre-trained model was fine-tuned for supervised classification of sequences with regards to the specific target they were enriched to bind (**Fig 1C**).

**Fig 1.**
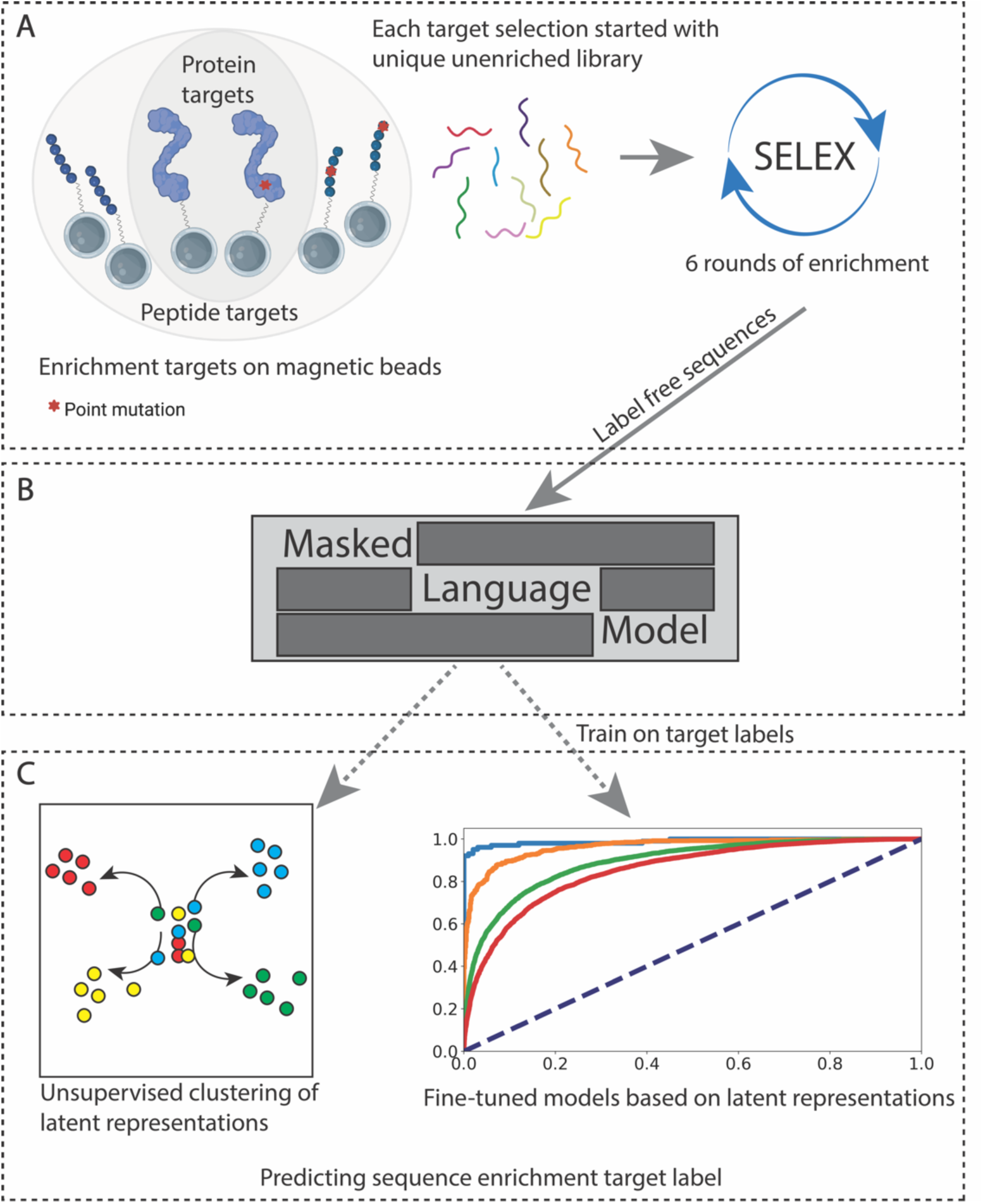
Overall analysis workflow. (A) Individual enrichments performed against protein and overlapping peptide targets, all started with unique unenriched libraries. (B) Label free sequences from selections used with a masked language model to create a pre-trained sequence model. This label free model can be used to (C) view unbiased sequence-based embeddings for trends in information content not trained on and to train predictive classifiers mapping target labels to ODN sequences

### Enrichment Targets

Two protein targets and four peptide targets spanning the wild-type or the mutant 2B (R1306Q) variant form of the Von Willebrand Factor (vWF), a blood glycoprotein involved in primary hemostasis, were used to explore the target specificity of latent space sequence representations created by a model based on natural language processing of unenriched and SELEX-enriched ssDNA libraries (**Table 1**). The recombinant vWF-A1-WT domain (207 aa) used in this study was previously used for SELEX development of a DNA aptamer (ARC1172)^42^. We also selected against the recombinant domain with a known point mutation (vWF-A1-2B: R1306Q) associated with increased bleeding due to platelet clearance from the blood^43,44^. In addition to the two protein domains, two overlapping 30 amino acid peptide fragments from each domain spanning the point mutation site were also used as enrichment targets. The peptides overlapped each other by 12 amino acids for both wildtype and mutant versions, representing varying degrees of sequence similarity, either sharing 40% within overlapping peptides or differing only by one residue between wildtype and mutant pairs. By comparing the latent representations of sequences from enriched libraries from these targets, and from unenriched starting libraries, the target-specificity of the information captured in the model latent space representations can be explored.

**Table 1.**
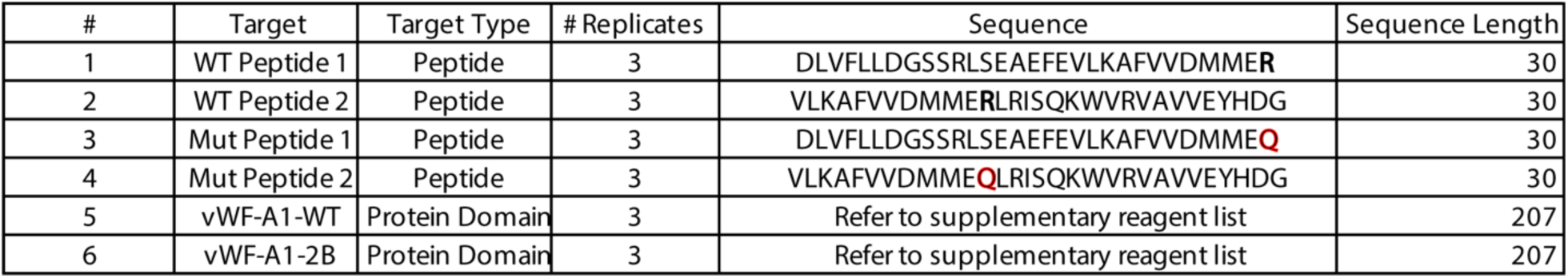
Target Peptides and Protein Domains Used.

For each target, the SELEX-enrichment was performed in triplicate, as described in Methods, starting with a unique unenriched library in each case. The 6 targets were conjugated to magnetic beads for the enrichment process and empty linker beads were used as control targets (beads with just the linker attached, but no target) to control for non-specific enrichment against the solid supports used. All targets were enriched together and under the same conditions on a high-throughput 96-well platform. A summary of sequence copy numbers from NGS results from the unenriched and round 6 enriched libraries are given in **S1 Table**.

### Binding of Enriched Sequences to Targets

Total binding of the enriched library pool for each peptide or protein target was determined by measuring the amount of bound biotinylated ssDNA to the bead-conjugated targets after incubation and washing (Methods). Quantitative determination of bound biotinylated library was performed using streptavidin-HRP detection measuring absorbance at 450 nm. The average absorption of three replicates is shown in **Fig 2**. As one might expect, the vWF domain and its point mutant gave statistically similar results. Somewhat more surprisingly, the library enriched using peptide 1 (WT Peptide 1) shows substantial binding, however the library enriched against mutant peptide 1 (Mut Peptide 1) showed binding levels comparable to the background binding of the unenriched library. Both wild type peptide 2 (WT peptide 2) and the corresponding point mutant (Mut Peptide 2) resulted in enriched libraries that bound significantly to their targets but Mut Peptide 2 enriched library bound significantly stronger to its target than WT Peptide 2 enriched library did its target. This difference was observed across all replicate enrichments as shown by the binding error bars in Fig 2. Thus, small changes in peptide but not protein sequences appear to result in changes in binding of the enriched libraries.

**Fig 2.**
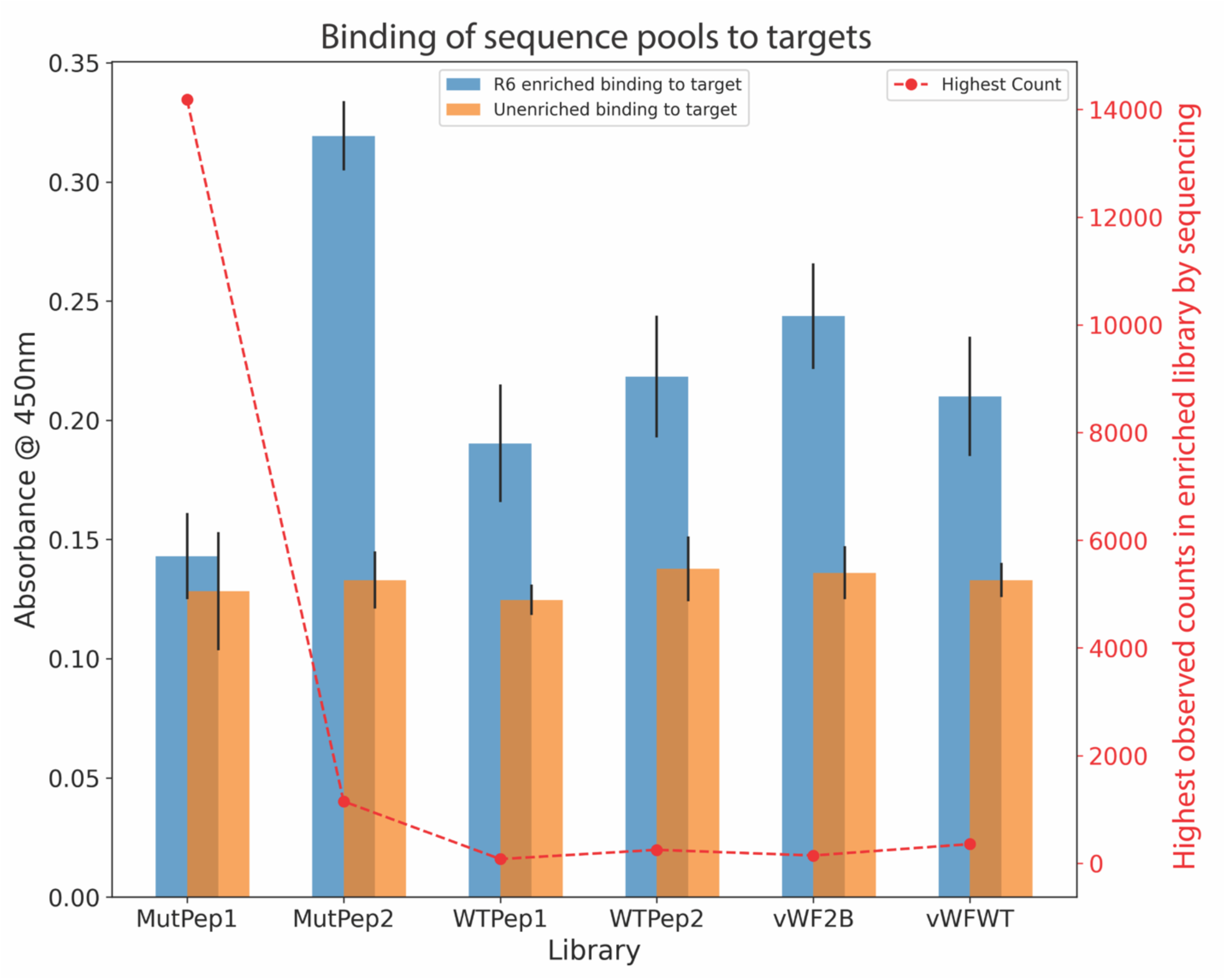
Differential Binding to Targets between Unenriched and Enriched Libraries. Absorption at 450nm was used to measure the binding of biotinylated DNA sequences to targets. Average binding across the three enrichment replicates to the targets plotted. Highest sequence copy number observed in libraries plotted on the secondary Y axis.

### Sequence based modeling

To identify patterns in the sequence data, we used a masked language model (MLM) architecture that consisted of three convolutional layers followed by two stacked biLSTM layers which were then concatenated and fed into a time distributed dense layer producing a per token prediction score (**S1 Fig**). The sequencing results were split into two files – training and holdout. The training file contained sequences randomly sampled from each sequence set and these were used to train the model in a self-supervised fashion without label information. The training file contained 500,000 sequences from each unenriched and enriched library except for three cases where there were too few sequences in those enriched libraries (specifically enriched libraries against Mutant peptide 1 replicate 2, from which we sampled 100,000 sequences and replicates 1 and 3 of the linker bead controls from which we were able to sample 400,000 sequences each). The sequences not used in the training were placed in the holdout set, which is never seen by the model, and used later for validation and model testing. In total, the training file consisted of 10.4 million sequences from across the different enrichments and the holdout file consisted of over 53 million sequences (**S2 Table)**. Thus, the MLM was trained on a set of sequences randomly sampled from each sequenced library and thus provided an overall view of enrichment rather than a target specific one. The self-supervised training was performed by masking 15% of the nucleotides across the sequences and then predicting these masked tokens and checking accuracy via backpropagation. This enabled the model to learn contextual information about the sequence space provided (this is referred to as the “pre-trained” model below to distinguish it from the fine-tuned models used for classification). The learned weights from the pre-trained model were then saved and used later to extract feature matrices for any input sequence which could then be used to generate latent space representations.

### Unsupervised analysis of the latent space sequence representations

**Fig 3** depicts a comparison of unsupervised clustering of three pairs of libraries. Dimension reduction was performed using Uniform Manifold Approximation and Projection (UMAP)^45^. For each comparison, tokenized sequences from the two sequence libraries being compared were merged and fed into the pre-trained model to generate latent space sequence representations (output embeddings/high-dimensional feature matrices. These output embeddings were then used as input to create 2-dimensional label free UMAP visualizations and then the sequence points were colored according to selection target label. Kernel Density Estimates (KDE) of distinct sequence sets were then calculated for each 2-dimensional representation to quantify the degree of overlap of the sequence latent space for different samples. The overlap in latent representations between two completely distinct unenriched libraries was calculated to be 95%, showing almost complete overlap. This high degree of overlap was also observed between three independent replicate enrichments against the same target with calculated KDE overlap of 87%. This high degree of overlap was observed between all replicate enrichments in the study (**S2 and S3 Figs**). Finally, latent representations of the library enriched against mutant peptide 2, which was observed to bind to its target, in comparison to unenriched sequences resulted in an overlap of only approximately 20%. What these results imply is that the latent space representations of the enriched sequences from mutant peptide 2 have captured information distinct to that target in comparison to unenriched sequences. This was done in a label free manner and across three technical replicates. However, latent space representations of enriched sequences resulting from enrichments starting with distinct random libraries as input, but all using mutant peptide 2 as a target, are nearly indistinguishable in terms of overlap between UMAP dimension reduced visualizations, like the comparison between two unenriched libraries. **Fig 4A** shows the KDE overlap results for all possible library comparisons using different targets. In general, overlap is very high between replicates (diagonal) but quite variable when comparing libraries from different targets. The variable KDE overlap between each target’s library and unenriched sequences, however, correlates strongly and inversely (Pearson correlation coefficient = -0.86, **Fig 4C**) with the binding (**Fig 1)**, implying that libraries that have the least overlap in sequence latent space representation with unenriched libraries also bind their targets most effectively. Thus, sequence latent space representations contain binding information. The error bars for overlap and binding are generated by comparing replicate enrichment sequence data for the targets. Additional visual comparisons of latent space representations between sequence sets from different targets or replicate targets are also provided (**S4 – S7 Figs**).

**Fig 3.**
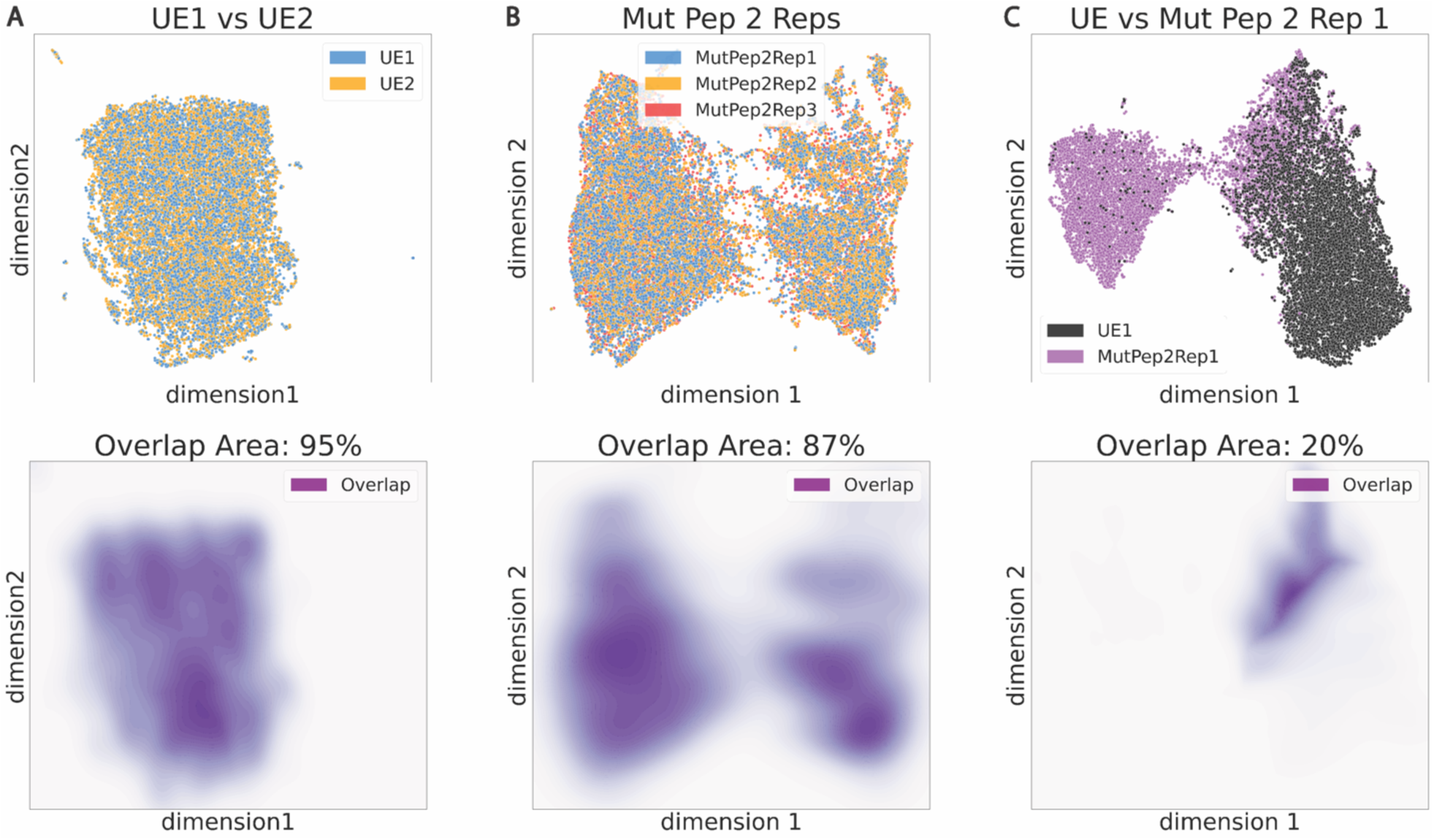
UMAP visualization of sequence latent space representations. (**Top Row**) Latent space representations comparing (**A**) two unenriched libraries, (**B**) three separate replicate enrichments against mutant peptide 2, and (**C**) unenriched vs enriched library against mutant peptide 2. (**Bottom Row**) Kernel density estimate (KDE) plots and overlaps provided below each comparison.

**Fig 4.**
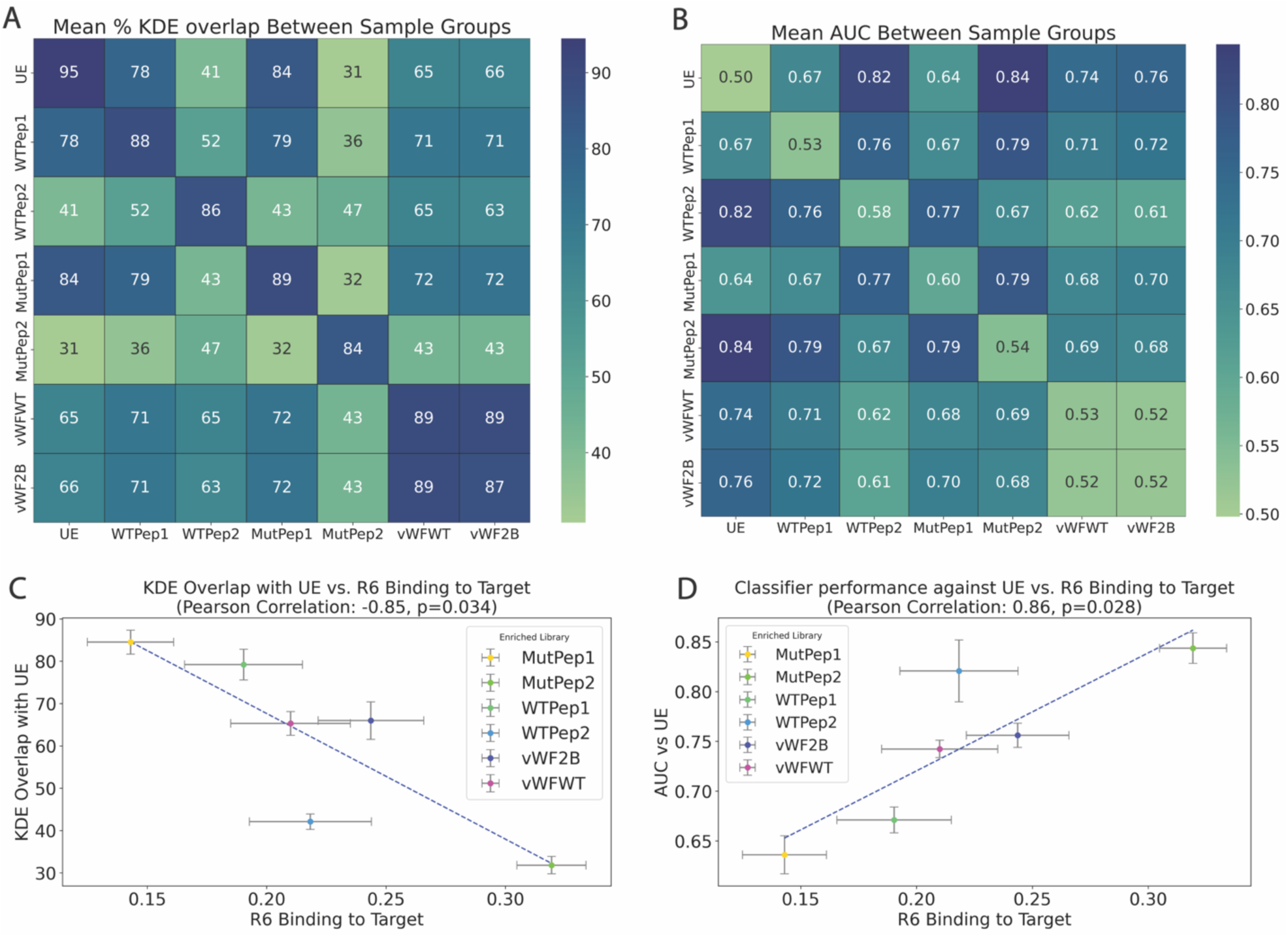
KDE overlap and AUC correlates with binding. **(A)** Mean KDE overlap values (%) between latent space representations of enriched pools comparing each possible pair of targets and with replicates on the diagonal. **(B)** Mean Area Under the Curve (AUC) from binary classification comparing sequences from each possible pair of targets and with replicates on the diagonal. **(C)** Linear fit of KDE overlaps vs binding for all possible comparisons between enriched targets and unenriched sequences. Standard deviations across the replicates were calculated and all found to be less than 6%. **(D)** Linear fit of AUC vs. binding for binary classification each enriched library vs. the unenriched library. Standard deviations across replicates were calculated with all comparisons less than 0.04. The highest standard deviation observed between replicate binding performance was 0.02

### Binary classification between sequence libraries using different targets

Another approach to determining the specificity of the models trained on enriched sequences for their targets is to perform binary classification between different target enriched libraries or between a target enriched library and an unenriched library. The pre-trained general model described above was modified by removing the time distributed dense layer from the original MLM architecture (S8 **Fig**) and then adding two dense hidden layers. For fine-tuning of the classifier, the weights from the MLM were inherited from the pre-trained model and fixed during the classifier training such that optimization occurs only on the newly learned weights from the two additional dense layers. Training between two sequence sets involved randomly selecting 50,000 sequences from each set without considering copy numbers obtained from sequencing, ensuring a diverse representation of each population. The sampled sequences were then split into training/test sets with an 80/20 ratio, respectively. Performance of the test set during the training was compared to performance of 50,000 holdout sequences using the same trained classifier. Classifier performances were reported as area under the curve (AUC) for the holdout set. We observed almost identical performance between test sequences and the holdout sequences, confirming that the model did not significantly overfit the data (**S3 Table**).

The results for all possible binary classification pairs using holdout sequences is depicted in Fig 4B. AUCs between unenriched libraries were 0.50 indicating no ability to classify the labels. This was also observed when using two simulated unenriched libraries (random sequences generated *in silico*) (**S9 Fig**). AUCs between target replicate enriched pools (diagonal) are also near 0.50. Classification between enriched samples is mixed, ranging from 0.52 to 0.84. AUC values between pairs are anticorrelated with KDE overlaps, as can be seen comparing **Fig 4A and Fig 4B**. A scatter plot comparing KDE overlap and classifier AUC values is given in **S10 Fig** and the Pearson correlation coefficient is -0.97. As with KDE overlap, AUC correlates strongly with binding when comparing enriched and unenriched libraries (**Fig. 4C and 4D**) with a Pearson correlation coefficient of 0.86 (compared to a -0.85 for KDE overlap). Thus, both KDE overlap and the AUC from binary classification with unenriched samples have the potential to serve as enrichment process optimization metrics. To assess the statistical significance of the classifier performances, permutation testing was performed. True labels in the classification were permuted 10,000 times and compared to classifier results to determine the number of times the permuted labels resulted in a classification performance either comparable or higher than non-permuted labels and plotted as a distribution. All classifications except unenriched vs unenriched comparisons were statistically different from random label comparisons, having p-values below 0.05 (**S11 Fig** for more details and examples) indicating that the observed classifier performances were not due to random chance.

The variation in AUC between enriched libraries also varies systematically with target similarity among the peptides. AUC values fall into three groups depending on the similarity of the targets of enrichment (**Fig 5)**. Replicates (complete similarity) have the lowest AUC values near 0.50. Single amino acid changes between target peptides move the value to about 0.67. When the 30 residue peptides overlap by 11-12 amino acids (18-19 amino acids different), the AUC ranges from 0.76-0.79.

**Fig 5.**
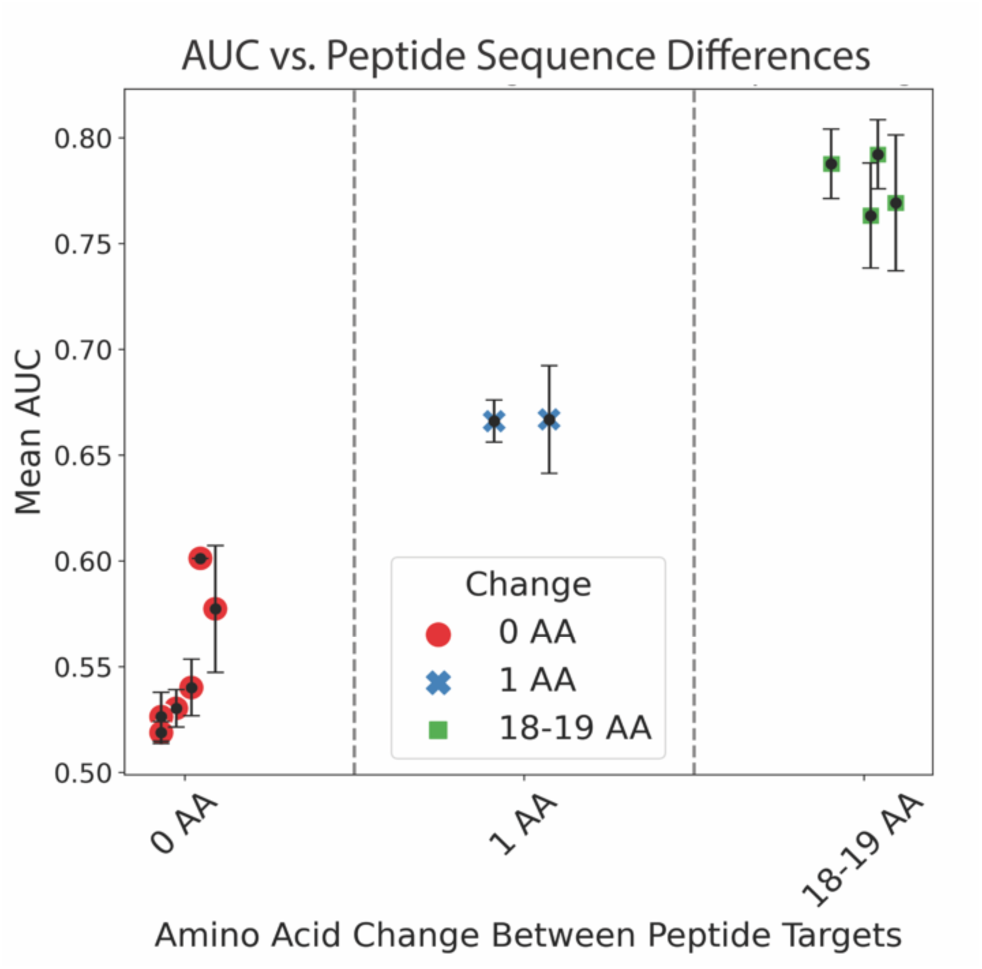
AUC values from classification between enriched libraries for three categories of sequence overlap between targets. Complete overlap, 1/30 amino acid changes, and 18-19/30 amino acid changes.

Thus, AUC values associated with binary classification correlate with KDE overlap, AUC also correlates well with binding when comparing enriched and unenriched sequence libraries and with the degree of similarity between peptides of the same length; both affinity and target specificity are captured in the latent space representations and the fine-tuned classifier models.

### Discriminating information content as a function of copy number in the library

Fine-tuned models were also trained as classifiers using sequences sampled based on the number of copies in the libraries enriched using mutant peptides 1 and 2 as targets. Mutant peptide 1 gives rise to an enriched library with very little binding while Mutant peptide 2 gives rise to a strongly binding enriched library (**Fig 1**). Samples compared included the sequences with the highest 10,000, 1000 and 100 copy numbers in the enriched sequence library and AUC values were determined relative to unenriched sequences. While there is effectively no consistent relationship between copy number and binding (**Fig 1, S1 Table**), an increase in AUC was observed with increasing copy number in mutant peptide 2 enriched sequences. However, there was no copy number dependence on classification with enrichments generated using mutant peptide 1 (**Fig 6**). The variable relationship between classification performance, copy numbers and binding was also observed in the other enrichments against the wildtype peptides (**S4 Table**). For example, while enriched pools against the vWF domains showed reasonable performance in classification from unenriched pools, the top 100 species by copy numbers from sequencing were found to not perform as well as the top 1k or 10k sequences. In fact, random sampling of sequences resulted in the best classification. Apparently, while the copy number of specific sequences is correlated with increased information content in some enrichment cases, this is not generally the case.

**Fig 6.**
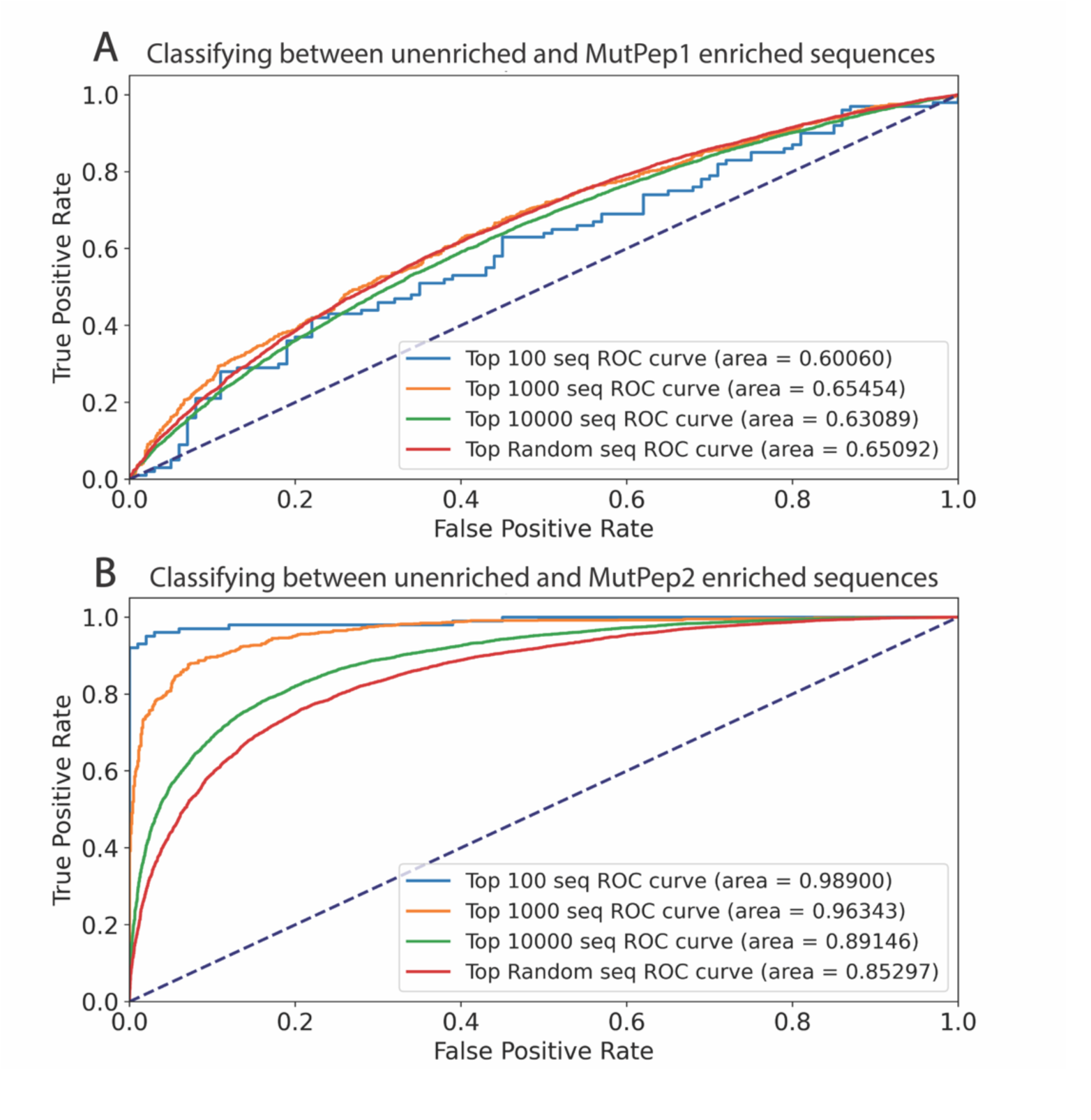
Receiver operator curves (ROC) resulting from variable sampling of sequences via copy numbers. **(A)** Classification comparing mutant peptide 2 enriched and unenriched sequences when sampling sequences via copy numbers. **(B)** Classification comparing mutant peptide 1 enriched and unenriched sequences when sampling sequences via copy numbers.

## Discussion

Enriched sequence libraries were created using 6 related target molecules: wild type and point mutant domains of vWF, two overlapping peptides from the wildtype domain and corresponding peptides containing the mutation. The binding of the enriched libraries was variable (**Fig. 2)**, with one of the mutant peptides giving rise to a library with little capacity to bind and the other peptides and the vWF wild type and mutant domains all directing moderate to high binding of the enriched library to the target. Sequences randomly sampled from across all the enriched libraries were combined and used to train a bidirectional Long Short-Term Memory (biLSTM) model using a Masked Language Model (MLM) approach. The resulting model weights were subsequently utilized both to generate latent space representations of sequences for unsupervised analysis and as the basis for fine tuning classifiers to distinguish enriched samples from each other or from unenriched libraries.

### Independent replicate enrichments resulted in essentially indistinguishable outcomes

Unsupervised clustering of latent space representations of sequences from independent enrichment replicates overlapped quantitatively (**Fig 3B and 4A**) indicating that the information content of the latent space representation is quite reproducible in enrichment to the same target starting from completely independent initial libraries with no sequence overlap. This is also reflected in the classification results comparing replicates to one another (**Fig 4B**); the enriched libraries were effectively indistinguishable even with supervised training. These results suggest that the latent space sequence representations impart recognition of structural similarities within enriched sequences that derive from independent, random starting sequence libraries.

### Binding is correlated with unsupervised clustering of latent space sequence representations and with classification accuracy

One of the most significant outcomes of this study is the finding that the binding is directly correlated with both the overlap of unsupervised clustering and the classification accuracy of supervised analysis between enriched and unenriched libraries (**Fig 4C and D**). Clustering overlap and classification accuracy are also highly correlated (**S10 Fig**). The implication of this, borne out in the analysis of sequence samples not involved in the training of the original general model or the fine-tuned classifiers at all, is that one can predict whether a given set of sequences, at least in aggregate, is likely to show significant binding to a target compared to unenriched sequences strictly from analyzing the latent space sequence representations. This implies that computational protocols to predict the composition of higher binding sequence sets should be possible.

### Copy number is not a general metric for binding potential of an enriched library to its target

Variable levels of specific sequence enrichment as gauged by sequence copy number was observed in enriched libraries across the different targets (S1 Table). Cluster separation in latent space and high classification performance between enriched and unenriched pools (**Fig 4**) was more indicative of target binding than was sequence copy number in an enriched library.

### Specificity of binding is also captured in the latent space representations and classifier models

Binding affinity is only part of what one would hope to capture in a predictive enrichment model. Just as important is the ability to predict specificity of binding. As shown in **Fig 5**, there are statistically significant differences in the classifier accuracy in considering three different well defined target comparisons: replicates using the same peptide target, comparison between a peptide and a single mutant of that peptide and comparison between two peptides that overlap in sequence by 11-12 amino acids (differ by 18-19 amino acids). At least in these cases, there is discernable differentiation between sequences as similar as one amino acid change in 30, implying that this level of specificity can be embedded in the latent space representation and learned by subsequent fine tuning to train a classifier.

### Predictive power of sequence-based models for SELEX enrichments

The most interesting conclusion from this work, both from a fundamental and practical perspective, is that machine learning models trained on sequence sets that aggregate enriched sequences from multiple distinct targets can learn what constitutes an enriched sequence, at least within the context of that set of targets. This suggests that there are common structural constructs in enriched sequence libraries among multiple targets, in agreement with past work^17,18,39^. Furthermore, the model demonstrates potential for identifying and characterizing these motifs, for example through an iterative process of sequence design, model evaluation, sequence modification, and structural modeling of optimized sequence classes. Fundamentally, this paves the way for studies investigating the mechanisms of enrichment from an information and structural perspective. At a practical level, it has the potential to allow more effective initial library design and more rapid optimization of protocols to achieve desired affinity and specificity of target binding.

## Methods and Materials

All peptides were synthesized by LifeTein (Hillsborough, New Jersey, USA), with products purified by HPLC to guarantee >98% purity. The two recombinant domains were purchased from IPAtherapeutics (Victoria, BC, Canada). All custom oligonucleotides were synthesized and purchased from Integrated DNA Technologies (Coralville, Iowa, USA). Pools of unenriched sequences consisted of a random 35 nucleotide variable region flanked by 2 constant primer regions, a 20-nucleotide anti-sense primer binding region and a 15-nucleotide sense binding region. The anti-sense primer was functionalized with a dibenzocyclooctyne (DBCO) moiety separated from the primer region via a poly A tail (35n in length) and 2 internal 18 carbon spacers to reduce interaction between the sequences and the azide beads used as a solid support. The sequence format of the unenriched pools and the primers are provided in **S5 Table. S6 Table** lists the reagents and kits used in this work and catalog numbers.

### Immobilizing targets to the beads

#### Target immobilization

The targets (peptides and domains) were immobilized on magnetic azide beads purchased from Click Chemistry Tools (Scottsdale, Arizona, USA) via a DBCO-S-S-NHS ester linker purchased from Conju-Probe (San Diego, California, USA). The disulfide in the linker allowed release of the target and bound sequences upon reduction with TCEP during the selection process. For coupling, an NHS ester on the linker is reacted first with N-terminal primary amines on the targets by incubating 48 nmoles of target with 24 nmoles of acetonitrile reconstituted linker in 100 µL of 1X PBS at pH 7.2 at room temperature and shaking at 500 RPM for 4 hours. After the incubation, the ester–primary amine reaction was quenched by addition of 1M Tris-HCl to a final concentration of 0.1M and then cooling on ice for 30 min. The quenched solution was then added to PBS-washed azide beads followed by gentle mixing. The total volume was then increased to 200 µL using additional 1X PBS. This was then mixed on a shaker at 500 RPM overnight at 4 ℃. After incubation, the magnetic beads were removed with a magnet and the supernatant was stored for HPLC analysis. The target immobilized beads were washed 3 times with 1X PBS, 0.05% Tween-20 and then stored in 200 µL at 4℃.

HPLC analysis: All HPLC runs were performed using a Thermofisher Scientific (Waltham, MA, USA) Vanquish autosampler instrument using a 250 X 2.1 (mm) Acclaim™ Vanquish C18 column, with the column and column pre-heater set to 60 ℃.

The HPLC gradient used is depicted in **S7 Table**, and **S12 Fig** contains histograms showing target-linker conjugated peaks and absence of these peaks in the supernatant of immobilization reactions with azide beads.

### Selection workflows

#### Selection protocol

Selection workflows were carried out using a KingFisher Flex bead processor from ThermoFisher Scientific (Waltham, MA, USA). All KingFisher Flex instrument protocol files can be provided upon request. The sample plates (unenriched single stranded DNA library in 1X PBS, 5mM MgCl_2_, 0.05% Tween-20), wash solutions (1X PBS, 5 mM MgCl_2_, 0.05% Tween-20), and elution solution (10 mM TCEP) were all loaded on the instrument prior to running. The target loaded beads were incubated with 200 ng ssDNA library for 1 hour to start Round 1 (4.0 x 10^12 unique species), followed by three washes for three minutes each and eluted at room temperature for 10 minutes. Each progressive round of selection saw a 10% reduction in starting library yield used and a one-minute increase in the mixing time of the beads per wash step. In round 4 two additional wash plates were introduced to increase selection stringency. The entire elution volume was used in the PCR reaction described after selection.

#### Sample preparation for SELEX round

The starting library was mixed with the required amount of 10X PBS (1X final), 1M MgCl2 (final 5mM) and water, mixed well and then heated to 95 ℃ for 5 minutes followed by rapid cooling to 4 ℃ for 10 minutes. This was then mixed with yeast tRNA and sheared salmon sperm DNA as competitor at a final concentration of 125 ng/µL each, along with pluronic F-127, a non-ionic surfactant, at a final concentration of 0.05%.

#### PCR amplification

PCR amplification was carried out using Q5 Hi fidelity hot start Taq-Polymerase (New England Biosciences) and the sample buffer provided. The PCR mix was incubated for 30 seconds at 98 ℃, followed by cycling as follows: 30 seconds at 98 ℃, followed by 30 seconds at 60 ℃, followed by 1 minute at 72 ℃. After all cycles, the reaction was held at 72 ℃ for 3 minutes followed by holding at 4 ℃.

#### Biotin strand recovery

After the PCR was complete, magnetic azide beads were used to capture the DBCO-tagged double stranded amplified DNA, then subjected to denaturing conditions to release the biotinylated strand using a KingFisher system to perform the bead handling. Excess primers and reducing agent were first removed from the PCR reaction using silica beads and a chaotropic binding (NTI buffer) step followed by an ethanol-based wash (NT3 buffer). NTI buffer is a guanidinium isothiocyanate buffer that is used in DNA purification systems to promote DNA binding to silica supports. NT3 buffer is an ethanol-based buffer that helps with washing away excessive salts and other impurities from the sample after binding to silica support. Both buffers were obtained from Macherey-Nagel (Düren, Nordrhein-Wesfalen, Germany). The magnetic silica bead cleanup was followed by capturing the double stranded product with azide beads. The azide beads were first washed in 5X PBS, then incubated with cleaned double stranded DNA in the presence of 5X PBS at 42 ℃ overnight. This incubation was followed by two 5-minute washes in 1 mL water each followed by a 20 second wash in 3 mM NaOH at room temperature to wash away any non-specifically bound double stranded material. Then a 10-minute elution in 100 µL of 30 mM NaOH at 56 ℃ was performed. The elution was quenched with 10 µL of 300 mM HCl and incubated at 4 ℃ for 15 minutes with shaking at 500 RPM.

### Sequencing

Sequencing was carried out for all libraries together in a multiplexed fashion, each sequence set was tagged with a unique predefined index primer obtained from Illumina (San Diego, USA). Sequencing reactions were prepared following the NGS prep protocol provided with the Illumina 50 cycle sequencing kit and performed on a NextSeq 550 instrument (Illumina). The results from each enrichment were compared pairwise to check for repetitions of the same species within different enrichments to confirm no potential cross contamination between enriched libraries.

### Binding tests

Binding of biotinylated enriched libraries to targets on beads was quantified using the reaction of Streptavidin – Horse Radish Peroxidase (HRP) to the biotinylated libraries followed by reacting HRP with 3,3’,5,5’ tetramethylbenzidine (TMB) substrate. Libraries (50 ng) were incubated with 5 µL of target-loaded magnetic beads in 1X PBS, 0.05% Tween-20 for 30 minutes at 500 RPM, followed by washing 3X with 1X PBS, 0.05% Tween-20 for 1 minute each at room temperature shaking at 500 RPM. The washed library bound beads were incubated at room temperature with streptavidin-HRP for 15 minutes followed by 3X washing using the same mixing conditions. Then the washed beads were incubated in TMB substrate for 15 minutes followed by quenching by addition of sulfuric acid to a 0.1M final concentration. Absorbance was then read at 450nm.

### Sequence model

Sequencing results from all samples in this study comprised approximately 66.5 million unique sequences. For training purposes, we only used sequences of 35n length. All sequences shorter than 35n were dropped and all sequences over that length were trimmed. The training dataset was assembled comprised of randomly sampled sequences from each enrichment target consisting of a total of 10.4 million sequences (S2 Table). Additionally, a holdout dataset was also created consisting of the remaining ∼ 53 million sequences. The training dataset was used to train a Masked Language Model (MLM), using only sequence information, with no copy number data. The architecture used for the pre-training step is provided in S1 Fig. Fifteen percent of the nucleotides across the tokenized sequences were masked and the MLM was trained to predict the masked tokens. This enabled self-supervised contextual learning about the sequence space provided. The model was built using tensorflow and keras libraries in python. The MLM used a biLSTM architecture, utilizing 256 LSTM units and the pre-training itself was performed for 500 epochs. The model architecture and weights were saved and used both to create latent space sequence representations for UMAP analysis and to serve as the functional model in the fine-tuning process as well.

#### Creating UMAP visualizations

The pre-trained model was used to create latent space representations of the sequences. These vectorized representations were used as the basis for dimension reduction using the UMAP package in python (parameters: n_components = 2, n_neighbor = 15, min_dist = 0.1 and used the ‘correlation’ metric). The resulting UMAP output, 2 dimensional coordinates (n_components), were plotted using the Seaborn and Matplotlib libraries in python.

#### Fine-tuning models on target labels/Testing trained classifiers on holdout data

Classifier training/fine-tuning was performed using the keras library in python. To train a classifier, the weights from the MLM were inherited from the pre-trained model and fixed such that the only newly learned weights were from the last two additional dense layers (**S8 Fig**). An 80:20 train/test split was used for training. All possible unique classifier combinations between the 20 sequence sets (190 unique pairs) were trained. Fifty-thousand sequences sampled randomly from each label in the holdout set was used to validate the generalization capabilities of the fine-tuned models by comparing performance to the test sequences. The classifier performances were recorded as area under the curve (AUC). All reported AUCs in this study are with holdout sequences. The original labels were permuted 10,000 times and these new permuted distributions were compared to the predicted distributions of the original labels to obtain a p-value by calculating the number of permuted distributions that had an AUC equal or greater to the observed (unpermuted) AUC:

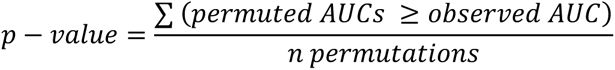

## Acknowledgments

We would like to thank Tassilo Hornung for his immense contributions in the early stages of development of computational workflows used in this study. We are very thankful for insightful discussions regarding this project with Michael Famulok and Günter Mayer. V.M is thankful for productive conversations with Ravi Chakra Turaga during analyses and writing. V.M is also very thankful to Caris Life Sciences for overall support throughout the undertaking of this study.

## Supporting information

**Table S1.** Results for 22 sequence sets are displayed showing total number of sequences, total number of unique sequences, and the results for the numbers of species at or above certain numbers.

| Sample | Species with counts |  |  |  |  |  |  |
| --- | --- | --- | --- | --- | --- | --- | --- |
|  | >=1 | 1 | >1 | >50 | >100 | >500 | >1000 |
| UE1 | 8.29E+06 | 8.20E+06 | 9.18E+04 | 0 | 0 | 0 | 0 |
| UE2 | 1.20E+07 | 1.18E+07 | 1.48E+05 | 0 | 0 | 0 | 0 |
| WT Peptide 1 - 1 | 1.80E+06 | 1.39E+06 | 4.13E+05 | 0 | 0 | 0 | 0 |
| WT Peptide 1 - 2 | 4.20E+06 | 2.21E+06 | 1.99E+06 | 81 | 5 | 0 | 0 |
| WT Peptide 1 - 3 | 4.81E+06 | 2.39E+06 | 2.41E+06 | 185 | 19 | 0 | 0 |
| WT Peptide 2 - 1 | 2.33E+06 | 1.17E+06 | 1.16E+06 | 14262 | 901 | 0 | 0 |
| WT Peptide 2 - 2 | 6.27E+06 | 3.70E+06 | 2.57E+06 | 0 | 0 | 0 | 0 |
| WT Peptide 2 - 3 | 4.71E+06 | 2.01E+06 | 2.69E+06 | 3 | 0 | 0 | 0 |
| Mut Peptide 1-1 | 5.65E+05 | 3.80E+05 | 1.85E+05 | 33113 | 21597 | 5196 | 1703 |
| Mut Peptide 1-2 | 1.04E+05 | 6.11E+04 | 4.30E+04 | 5554 | 2186 | 17 | 1 |
| Mut Peptide 1-3 | 8.77E+05 | 5.55E+05 | 3.22E+05 | 54298 | 31642 | 2357 | 129 |
| Mut Peptide 2-1 | 2.84E+06 | 1.36E+06 | 1.48E+06 | 12089 | 2378 | 15 | 0 |
| Mut Peptide 2-2 | 2.71E+06 | 1.32E+06 | 1.39E+06 | 12804 | 2964 | 34 | 2 |
| Mut Peptide 2-3 | 1.65E+06 | 8.21E+05 | 8.25E+05 | 31274 | 9224 | 168 | 12 |
| vWF_WT-1 | 2.50E+06 | 1.12E+06 | 1.37E+06 | 2188 | 44 | 0 | 0 |
| vWF_WT-2 | 2.46E+06 | 1.06E+06 | 1.39E+06 | 1464 | 16 | 0 | 0 |
| vWF_WT-3 | 1.51E+06 | 8.09E+05 | 7.03E+05 | 32476 | 3765 | 0 | 0 |
| vWF_2B-1 | 1.56E+06 | 7.36E+05 | 8.20E+05 | 196 | 3 | 0 | 0 |
| vWF_2B-2 | 1.90E+06 | 8.56E+05 | 1.04E+06 | 2602 | 84 | 0 | 0 |
| vWF_2B-3 | 2.51E+06 | 1.10E+06 | 1.41E+06 | 2247 | 43 | 0 | 0 |
| LBeads-1 | 4.92E+05 | 3.29E+05 | 1.63E+05 | 42443 | 31096 | 1624 | 73 |
| LBeads-3 | 4.40E+05 | 3.05E+05 | 1.35E+05 | 26799 | 23120 | 8050 | 1845 |

**Table S2.** Number of sequences used in training the pre-trained model and the number of sequences remaining in the holdout sets.

| Sequence Set | # sequences in train | # sequences in holdout |
| --- | --- | --- |
| WT pep 1 -1 | 5.00E+05 | 1.27E+06 |
| WT pep 1 -2 | 5.00E+05 | 3.49E+06 |
| WT pep 1 -3 | 5.00E+05 | 4.06E+06 |
| WT pep 2 -1 | 5.00E+05 | 1.66E+06 |
| WT pep 2 -2 | 5.00E+05 | 5.58E+06 |
| WT pep 2 -3 | 5.00E+05 | 3.96E+06 |
| Mut pep 1 -1 | 5.00E+05 | 5.87E+04 |
| Mut pep 1 -2 | 1.00E+05 | 3.83E+03 |
| Mut pep 1 -3 | 5.00E+05 | 3.46E+05 |
| Mut pep 2 -1 | 5.00E+05 | 2.14E+06 |
| Mut pep 2 -2 | 5.00E+05 | 2.06E+06 |
| Mut pep 2 -3 | 5.00E+05 | 1.02E+06 |
| vWF-WT -1 | 5.00E+05 | 1.81E+06 |
| vWF-WT -2 | 5.00E+05 | 1.81E+06 |
| vWF-WT -3 | 5.00E+05 | 9.02E+05 |
| vWF-2B -1 | 5.00E+05 | 1.03E+06 |
| vWF-2B -2 | 5.00E+05 | 1.33E+06 |
| vWF-2B -3 | 5.00E+05 | 1.83E+06 |
| Lbeads -1 | 4.00E+05 | 8.57E+04 |
| Lbeads -3 | 4.00E+05 | 3.39E+04 |
| UE-1 | 5.00E+05 | 7.75E+06 |
| UE-2 | 5.00E+05 | 1.14E+07 |
| Total | 1.04E+07 | 5.36E+07 |

**Table S3.**
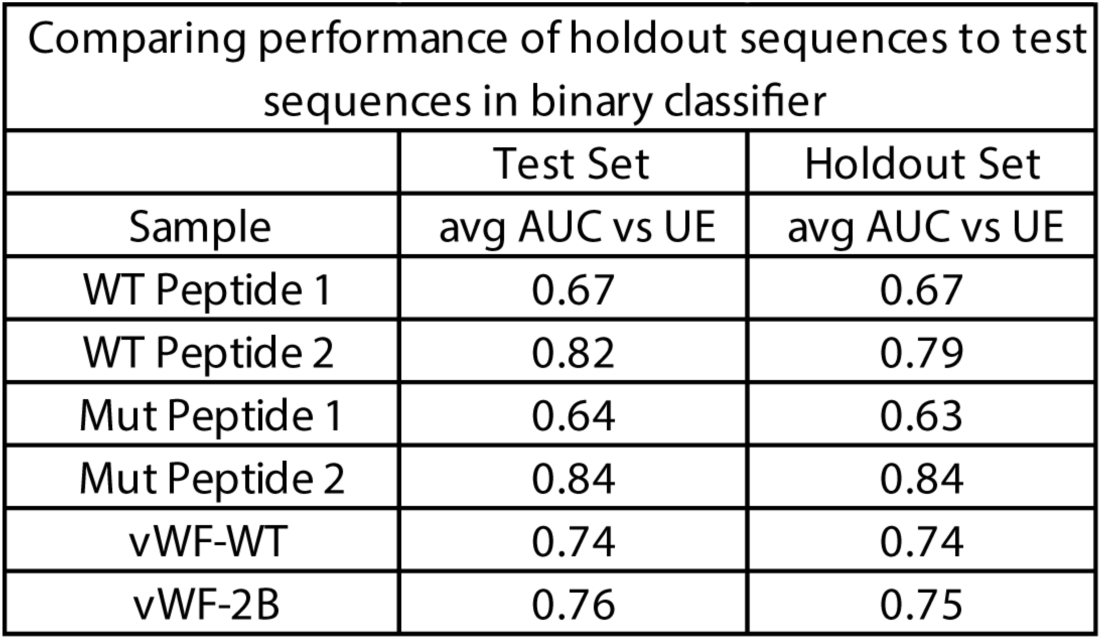
Classifier performances reported as AUCs between test and holdout sequences. Main results are reported using holdout sequences.

| Comparing performance of holdout sequences to test sequences in binary classifier |  |  |
| --- | --- | --- |
|  | Test Set | Holdout Set |
| Sample | avg AUC vs UE | avg AUC vs UE |
| WT Peptide 1 | 0.67 | 0.67 |
| WT Peptide 2 | 0.82 | 0.79 |
| Mut Peptide 1 | 0.64 | 0.63 |
| Mut Peptide 2 | 0.84 | 0.84 |
| vWF-WT | 0.74 | 0.74 |
| vWF-2B | 0.76 | 0.75 |

**Table S4.** Variable classifier performance observed with sequence sampling via copy numbers across the different enrichments. Variable performances were similar between replicates.

|  | AUC vs unenriched |  |  |  |
| --- | --- | --- | --- | --- |
| Target | Random | Top 10k | Top 1k | Top 100 |
| WT Peptide 1 | 0.67 | 0.71 | 0.77 | 0.81 |
| WT Peptide 2 | 0.82 | 0.85 | 0.92 | 0.95 |
| Mut Peptide 1 | 0.64 | 0.64 | 0.66 | 0.66 |
| Mut Peptide 2 | 0.84 | 0.89 | 0.97 | 1.00 |
| vWF-A1-WT | 0.74 | 0.74 | 0.72 | 0.69 |
| vWF-A1-2B | 0.76 | 0.76 | 0.72 | 0.68 |

**Table S5.**
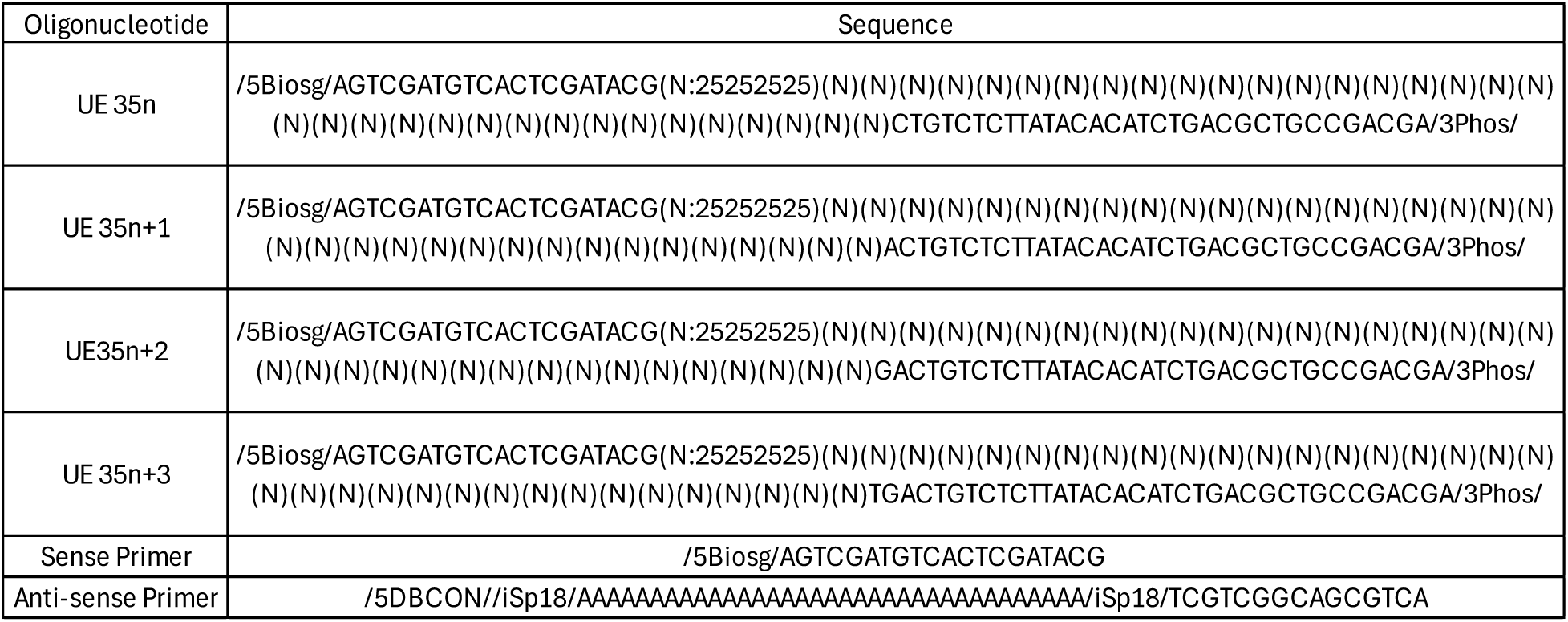
Oligonucleotide details. Oligonucleotide orders were placed through IDT (Integrated DNA Technologies – Coralville, IA) via Ultramer synthesis. The 35n region was designed to maintain an equal representation of the 4 nucleotides. Four different syntheses of unenriched sequences were mixed in equimolar amounts to form the master unenriched library used in the study.

**Table S6.** List of reagents and consumables used in experimental workflows with supplier and catalog information provided.

| Product | Supplier | Catalog # |
| --- | --- | --- |
| 1M Tris HCl, pH 8.0 | Thermofisher Scientific | 15586-025 |
| PBS Salt | Sigma-Aldrich | P3813 |
| 10% Pluronic F-127 | Thermofisher Scientific | P6866 |
| 10% Tween-20 | Sigma-Aldrich | P9416 |
| 1M MgCl<sub>2</sub> | Thermofisher Scientific | AM9530G |
| Acetonitrile | Fisher chemical | A461-4 |
| Trifluoroacetic acid | Thermofisher Scientific | 28904 |
| 0.5M TCEP | Thermofisher Scientific | 77720 |
| Sheared Salmon Sperm DNA | Thermofisher Scientific | 1563-011 |
| Yeast tRNA | Thermofisher Scientific | AM7119 |
| Q5 Hi-Fidelity Hot Start PCR Kit | NEB | M0493L |
| 100mM dNTPs | Thermofisher Scientific | 10297117 |
| 10M NaOH | Sigma-Aldrich | 72068 |
| Sulfuric acid | Thermofisher Scientific | 039669.K2 |
| Formic acid | Sigma-Aldrich | 33015 |
| TMB Substrate | Thermofisher Scientific | 34028 |
| 4% EX Agarose Pre-cast gel | Thermofisher Scientific | G401004 |
| Azide beads | Click-Chemistry Tools | 1036-5 |
| Silica beads (PureBind M Beads) | Ocean-nanotech | P0202-050 |
| DBCO-SS-NHS ester linker | Conju-Probe | CP-2024 |
| NTI buffer | Macherey-Nagel | 740305.120 |
| NTC buffer | Macherey-Nagel | 740654.100 |
| NT3 buffer | Macherey-Nagel | 740598 |
| Qubit DS DNA HS kit | Thermofisher Scientific | Q32854 |
| Sequencing kit (50 cycle, SBS) | Illumina | 20024906 |
| NextSeq Flowcell | Illumina | 2022408 |
| 250 X 2.1 mm Acclaim Vanquish C18 column | Thermofisher Scientific | 074812-V |

**Table S7.** HPLC gradient to test target – linker conjugations and bead immobilizations. Buffer A: 99.9% H_2_O, 0.1% Formic acid, 0.02% Trifluoroacetic acid Buffer B: 99.9% acetonitrile, 0.1% Formic acid, 0.02% Trifluoroacetic acid

| Time (min) | Flow (mL/min) | %A | %B |
| --- | --- | --- | --- |
| 0 | 0.4 | 95 | 5 |
| 3 | 0.4 | 95 | 5 |
| 13 | 0.4 | 65 | 35 |
| 33 | 0.4 | 55 | 45 |
| 36 | 0.4 | 20 | 80 |
| 37 | 0.4 | 0 | 100 |
| 40 | 0.4 | 0 | 100 |
| 41 | 0.4 | 95 | 5 |
| 46 | Stop Run |  |  |

**Fig S1.**
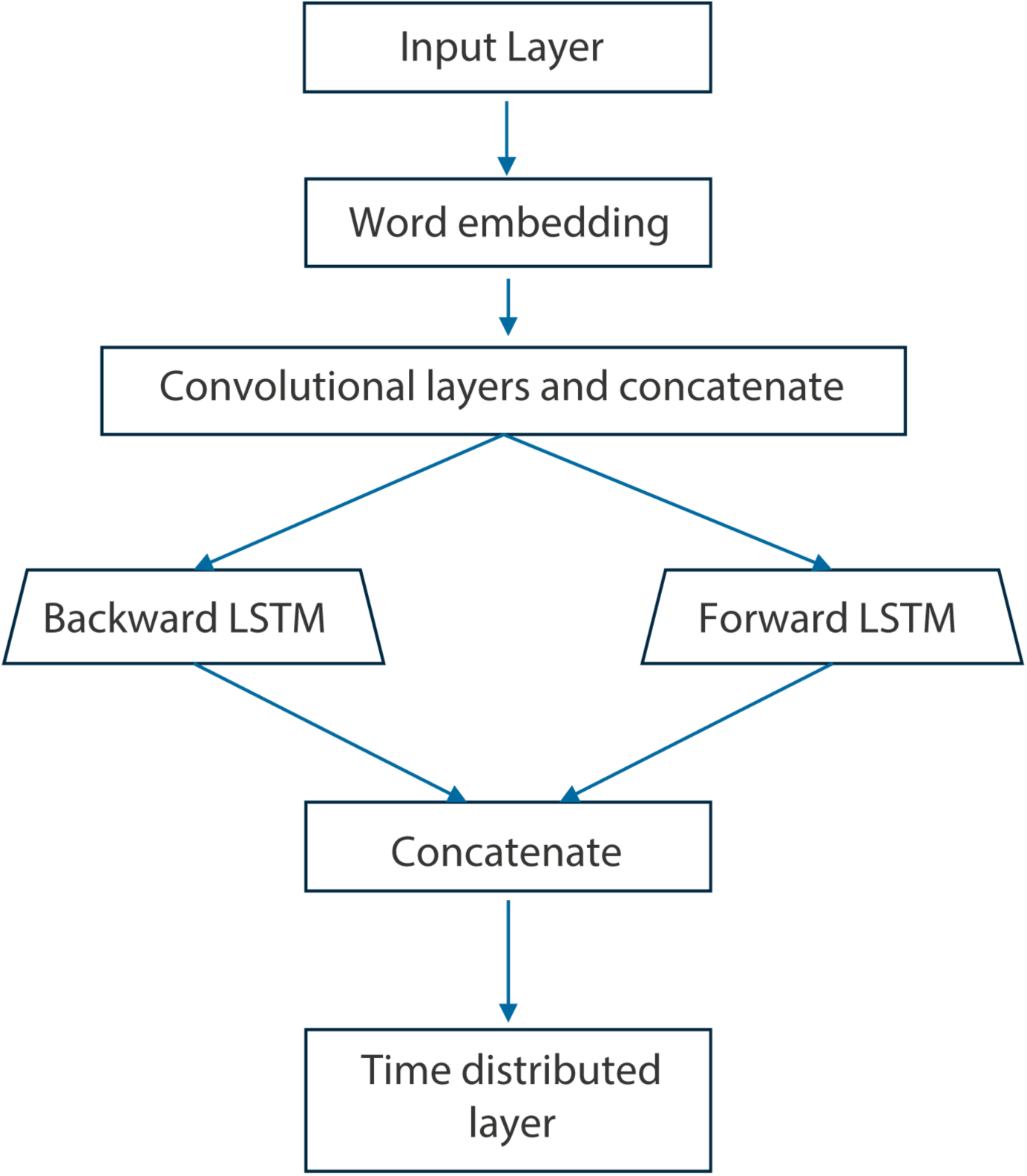
Simple illustration of pre-training architecture. Layer descriptions provided below: InputLayer – Sequence input layer Embedding – This layer converts input sequences into dense vectors of fixed size where each token in the sequence is represented as an embedded vector Convolutional layers: Used to capture local patterns in the data and then concatenated LSTM layers: Long-term sequence dependencies are captured using a biLSTM architecture with 256 units each Concatenate: LSTM layer outputs are merged to combine the learned features Time distributed layer: Fully connected dense layer to each time step, performing a classification on the learned features (semi-supervised)

**Fig S2.**
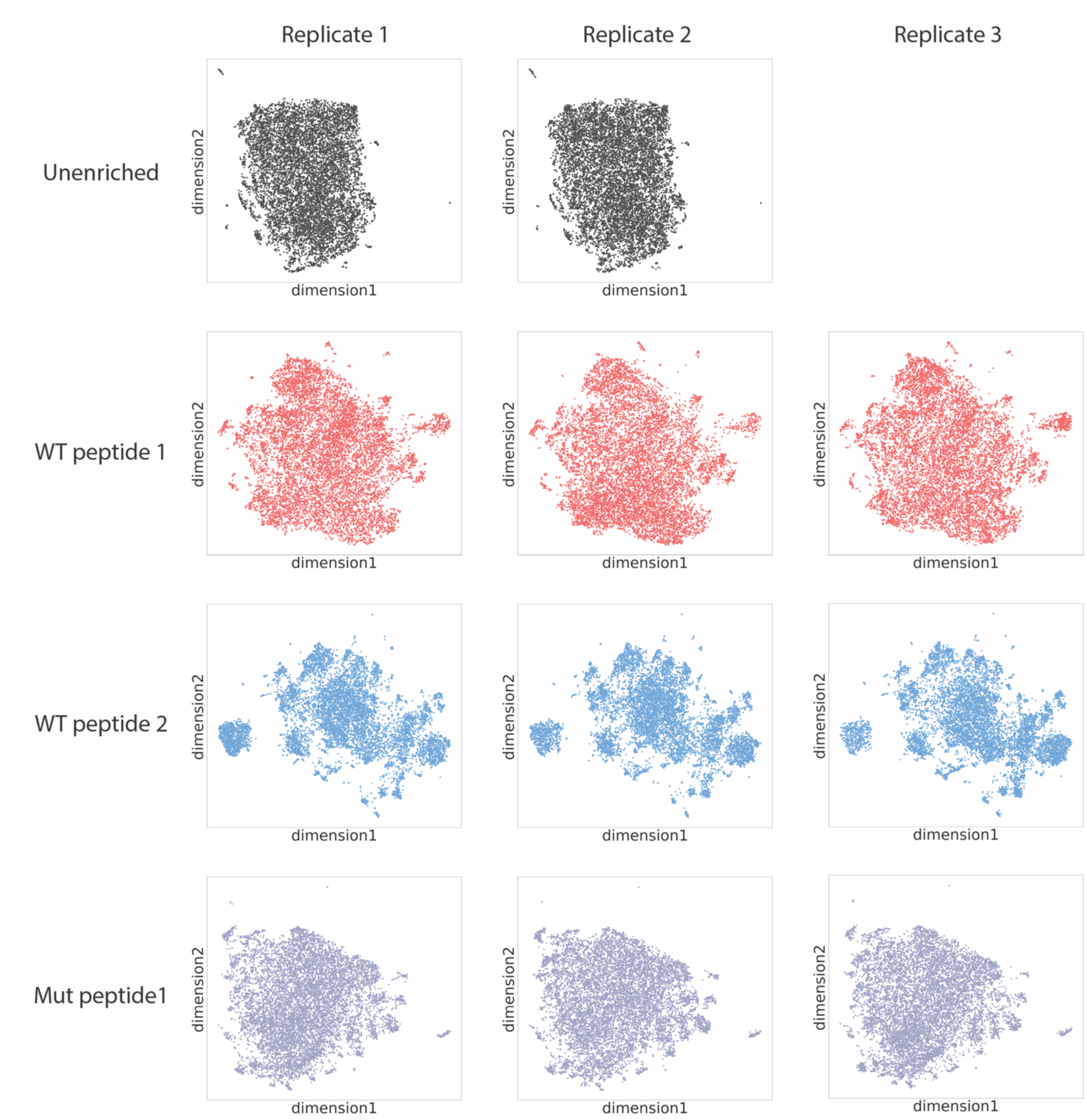
Similar UMAP representations between all enrichment replicate pools in latent space. These distributions in latent space are determined using the pre-trained model generated using label free sequences

**Fig S3.**
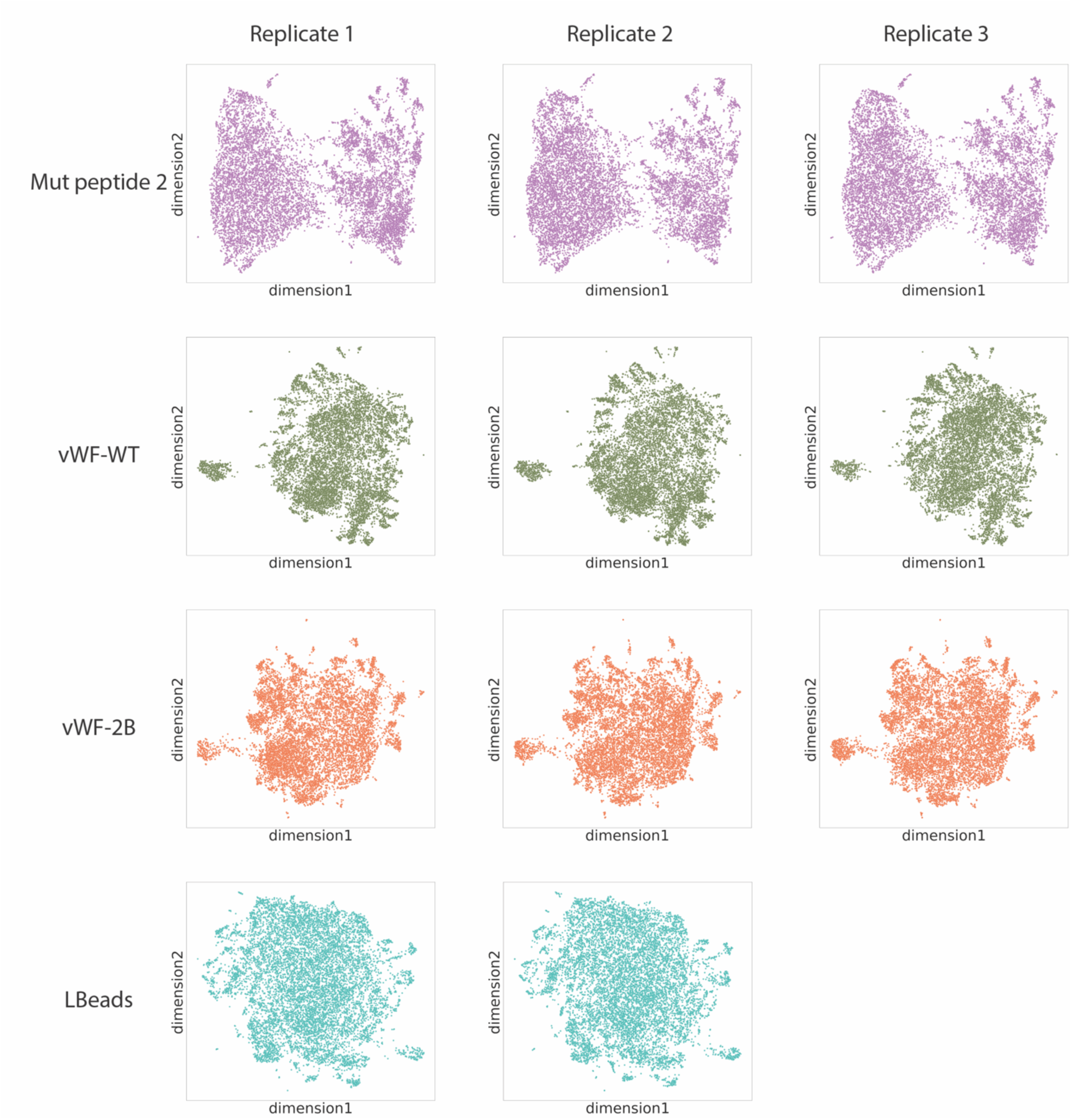
Similar UMAP representations between all enrichment replicate pools in latent space. These distributions in latent space are determined using the pre-trained model generated using label free sequences

**Fig S4.**
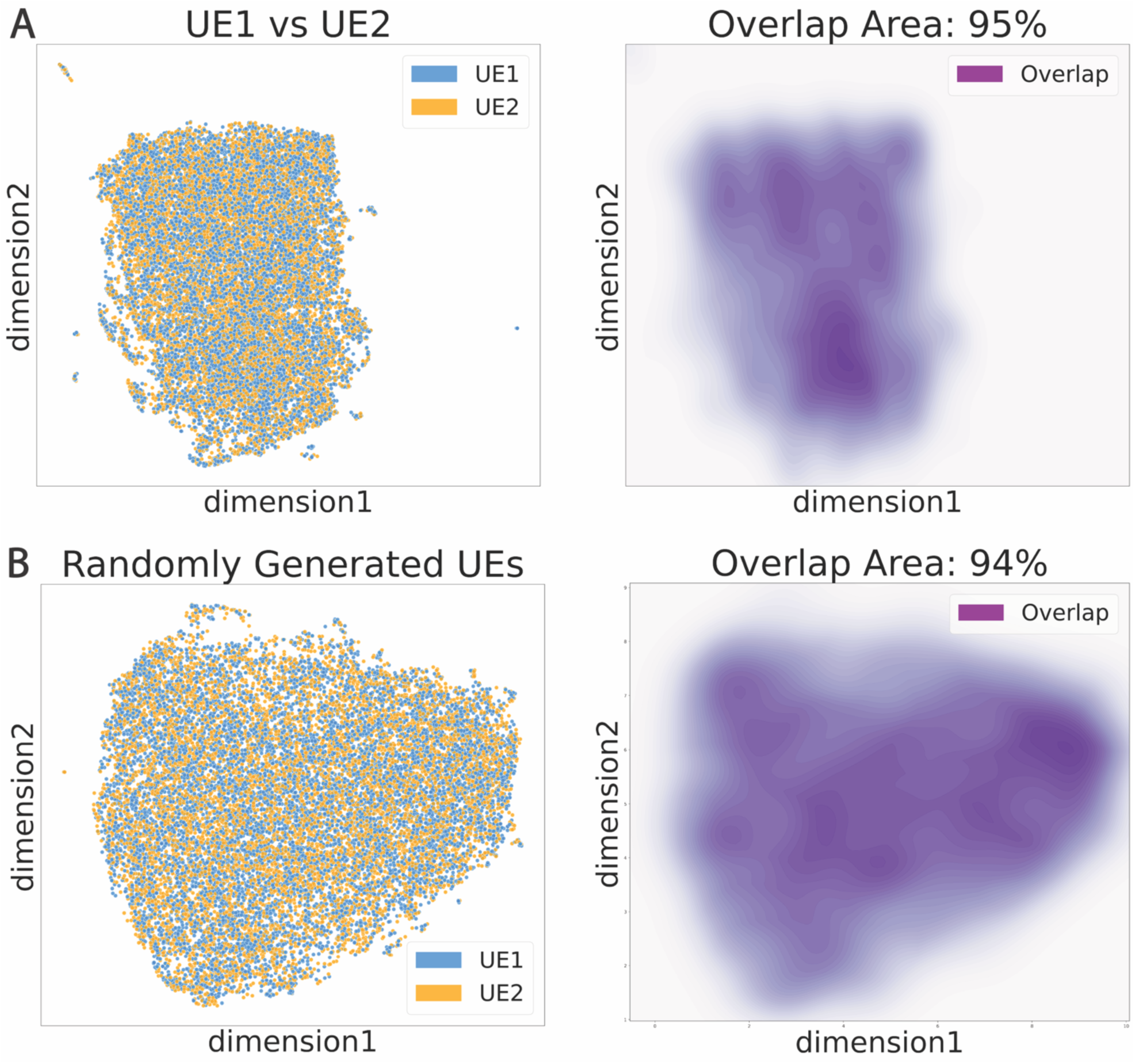
Latent space representations of unenriched sequences. Representations in the left column with KDE overlap plots on the right.

**Fig S5.**
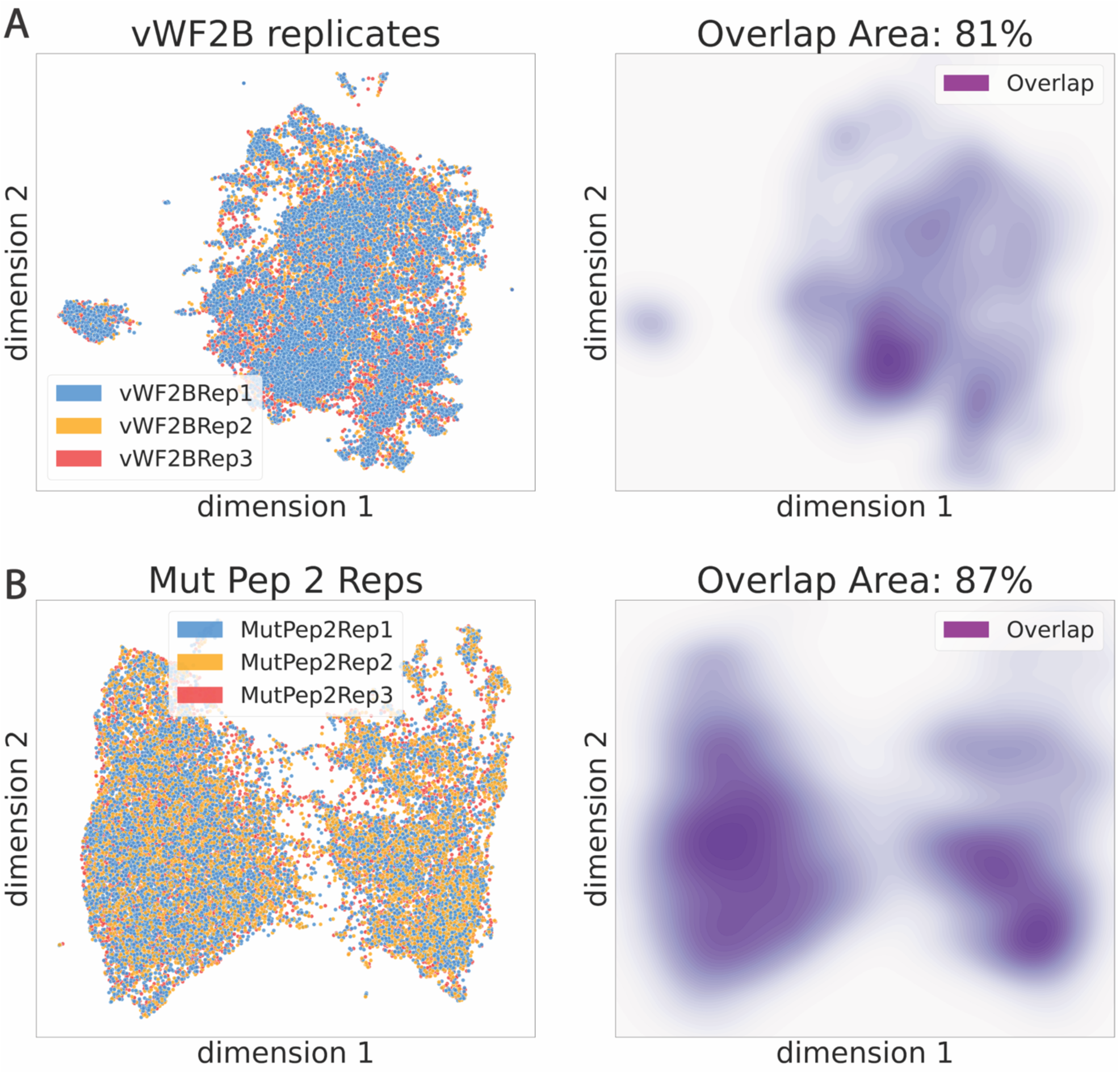
Latent space representations of replicate enrichments. Representations in the left column with KDE overlap plots on the right.

**Fig S6.**
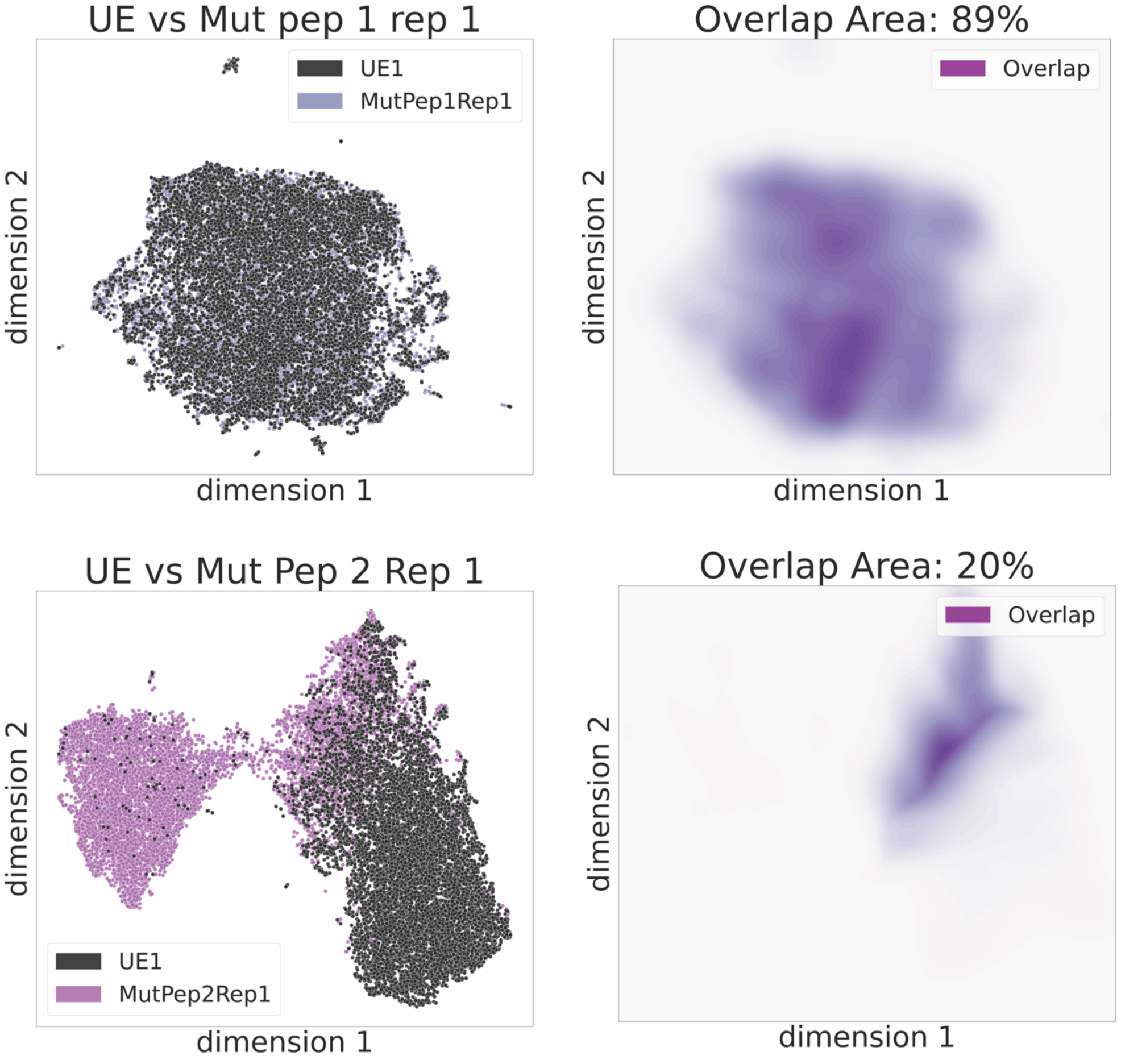
Latent space representations comparing unenriched and enriched sequences. Representations in the left column with KDE overlap plots on the right.

**Fig S7.**
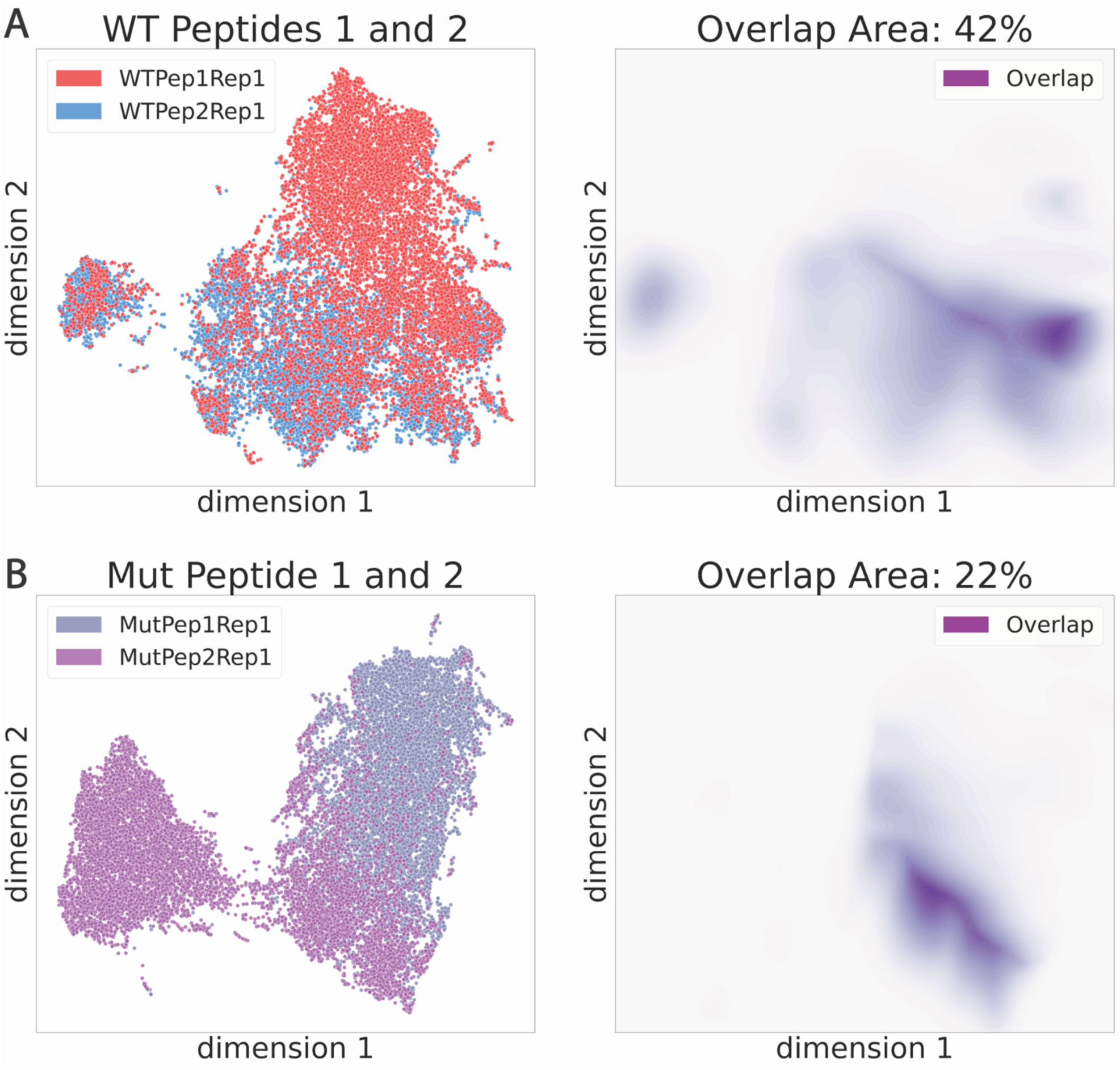
Latent space representations comparing enriched sequences. Representations in the left column with KDE overlap plots on the right.

**Fig S8.**
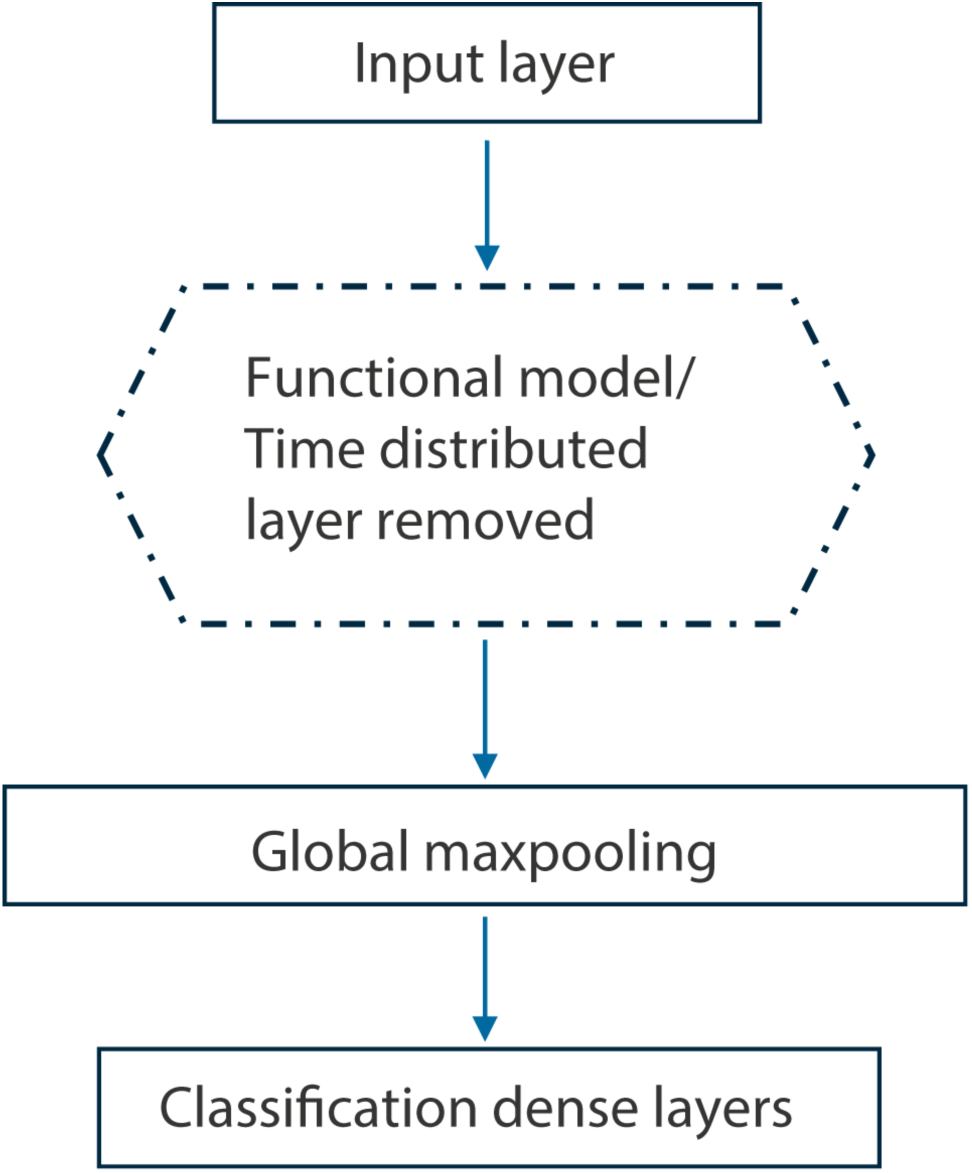
Fine-tuning architecture for training on selection target label. Layer descriptions provided below: InputLayer – Sequence input layer Functional – The entire pre-trained model architecture from S1 FigA with time distributed layer removed. Output is 512-dimensional feature map Global_max_pooling1d: Reduces feature map to its maximum value across entire sequence, distilling most prominent features from a sequence Dense layers: Fully connected layers responsible for binary classification

**Fig. S9.**
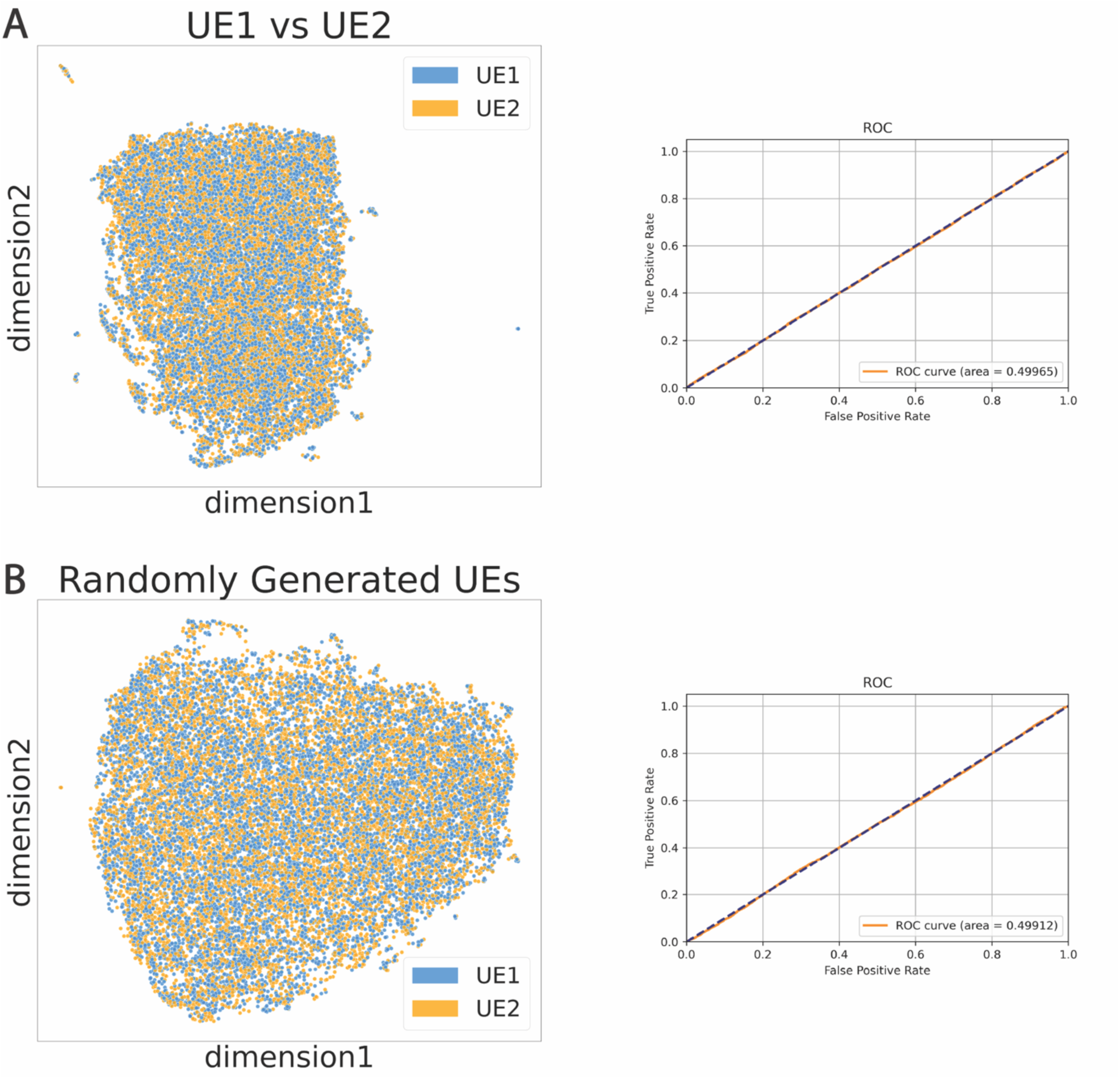
Latent space representations and classification accuracy comparisons between experimentally sequenced unenriched libraries and simulated unenriched libraries. There is no obvious unsupervised clustering in these examples and classification with fine-tuning is random.

**Fig S10.**
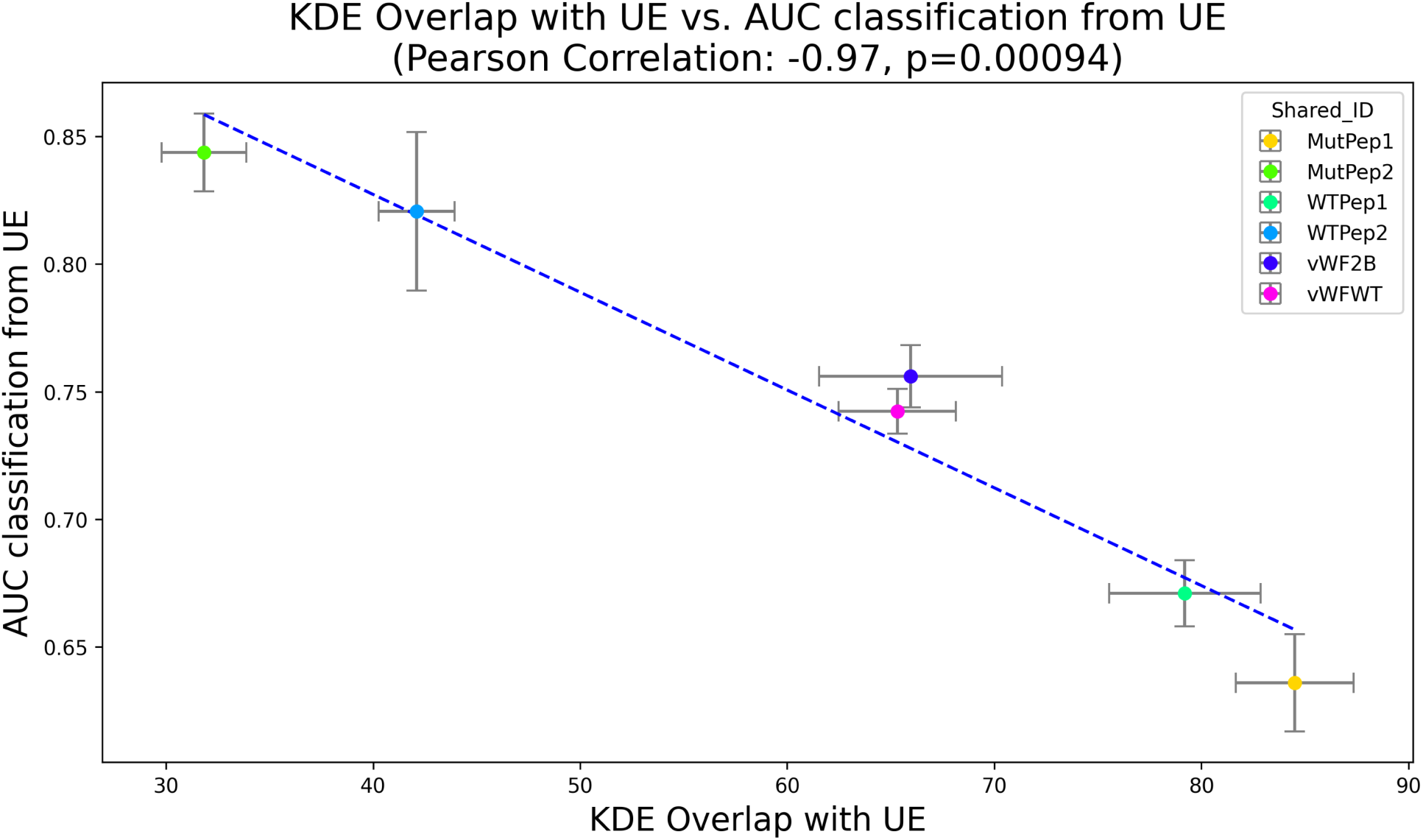
Linear relationship between KDE overlaps and classification AUC between sequence sets. Plot showing the linear relationship between KDE overlaps and classification AUCs when comparing unenriched and enriched sequence sets. The data was found to have an inverse but highly linear relationship, suggesting that classification accuracies can be inferred strongly using latent space overlaps between the sequence sets. The standard error points are derived from latent space overlaps and classification AUCs between unenriched and all three enrichment replicates.

**Fig S11.**
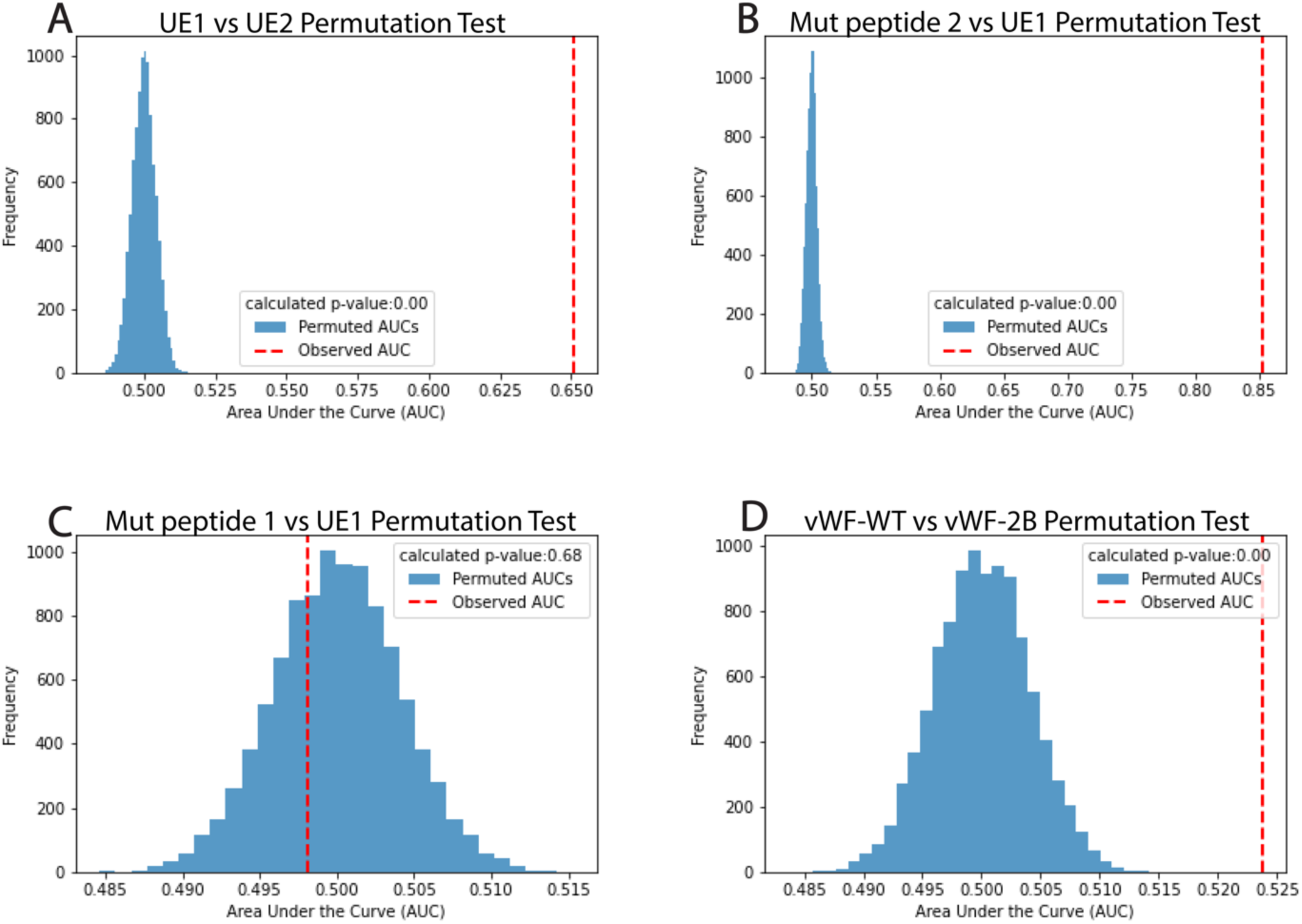
Permutation tests of the significance of trained classifier results. – **(A)** Permutation test resulted in a p-value of 0.68, indicating a result that can be obtained by chance, something expected for classifying between two unenriched sequence sets. With the introduction of target enriched libraries in classification vs unenriched sequences, the p-value drops below 0.05 for all classifications **(B-D)**, even in the case of enrichments that produced high-copy number libraries that did not bind to their targets, and that did not classify in comparisons with unenriched sequences.

**Fig S12.**
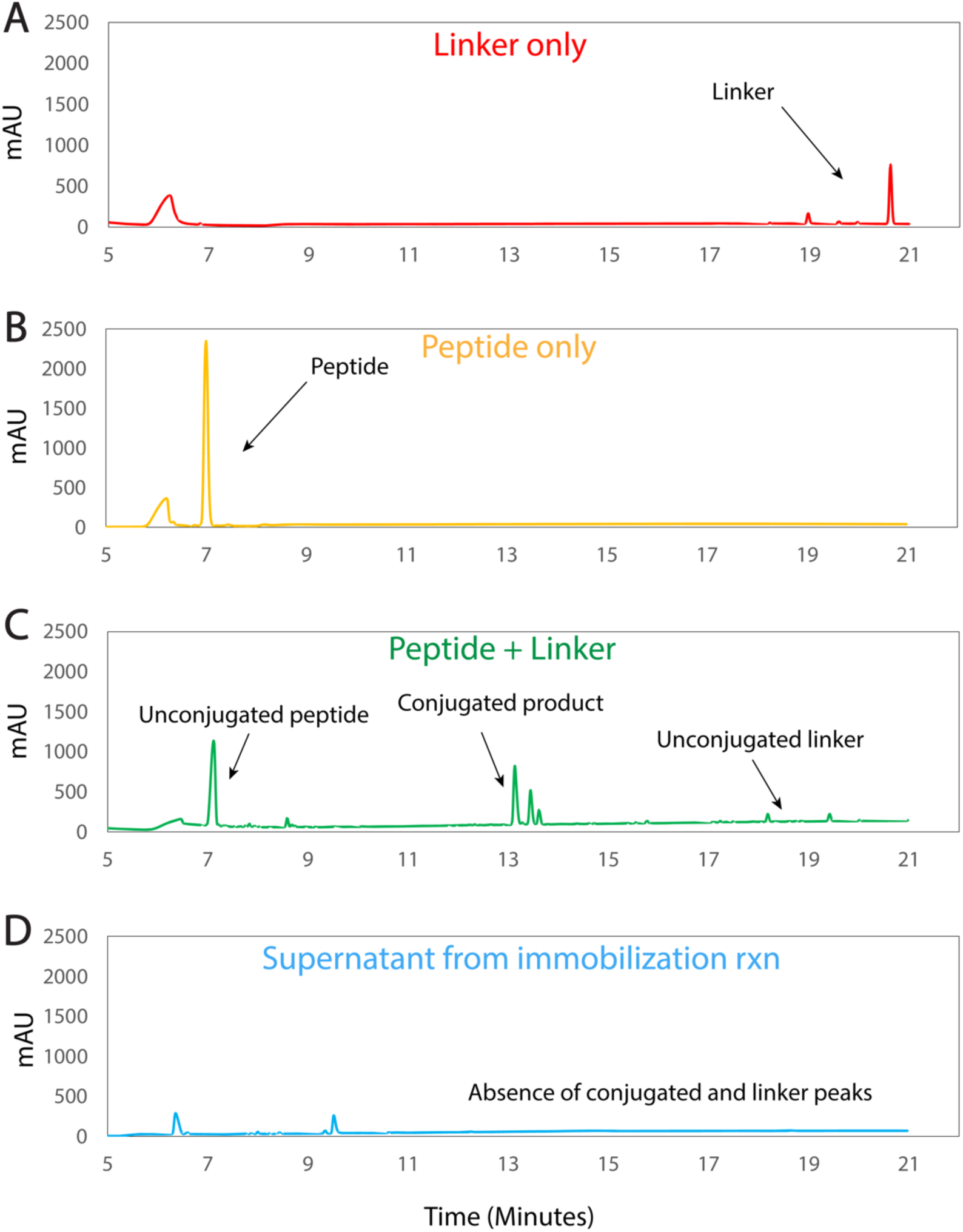
HPLC runs verifying conjugations and target immobilization to beads. HPLC chromatograms showing the absorbance of **(A)** Linker used for conjugations, **(B)** A peptide target by itself, **(C)** Conjugation reaction between a peptide target and linker showing unreacted linker and peptide with conjugated product in the middle and **(D)** Immobilization supernatant showing absence of conjugation and linker peaks confirming attachment to the beads.

## Author Contributions

**Conceptualization:** Varun Maher, Neal Woodbury, Daniel Martin, Heather O’Neill, David Spetzler

**Data curation:** Daniel Martin

**Formal analysis:** Varun Maher, Daniel Martin **Funding acquisition:** David Spetzler **Investigation:** Varun Maher, Daniel Martin **Methodology:** Varun Maher, Daniel Martin

**Project administration:** Heather O’Neill, Neal W. Woodbury.

**Resources:** Heather O’Neill, David Spetzler

**Software:** Daniel Martin

**Supervision:** Heather O’Neill, Neal W. Woodbury., Zhan-Gong Zhao

**Visualization:** Varun Maher

**Writing – original draft:** Varun Maher, Neal Woodbury

**Writing – review & editing:** Varun Maher, Neal Woodbury, Daniel Martin, Heather O’Neill

## Notes

### Competing Interest Statement

This work was carried out as a collaboration between Arizona State University and Caris Life Sciences. Personnel were paid by their respective institutions. All laboratory and computational work was performed at Caris Life Sciences using their resources.

## References

1. Tuerk, C. & Gold, L. Systematic evolution of ligands by exponential enrichment: RNA ligands to bacteriophage T4 DNA polymerase. Science 249, 505–510 (1990).

2. Ellington, A. D. & Szostak, J. W. In vitro selection of RNA molecules that bind specific ligands. Nature 346, 818–822 (1990).

3. Ellington, A. D. & Szostak, J. W. Selection in vitro of single-stranded DNA molecules that fold into specific ligand-binding structures. Nature 355, 850–852 (1992).

4. Lorsch, J. R. & Szostak, J. W. In vitro selection of RNA aptamers specific for cyanocobalamin. Biochemistry 33, 973–982 (1994).

5. Famulok, M. & Szostak, J. Stereospecific recognition of tryptophan agarose by in vitro selected RNA. J. Am. Chem. Soc. 114, 3990–3991 (1992).

6. Nieuwlandt, D., Wecker, M. & Gold, L. In Vitro Selection of RNA Ligands to Substance P. Biochemistry 34, 5651–5659 (1995).

7. Williams, K. P. et al. Bioactive and nuclease-resistant l-DNA ligand of vasopressin. Proc. Natl. Acad. Sci. 94, 11285–11290 (1997).

8. Chen, H., McBroom, D. G., Zhu, Y.-Q., Gold, L. & North, T. W. Inhibitory RNA Ligand to Reverse Transcriptase from Feline Immunodeficiency Virus. Biochemistry 35, 6923–6930 (1996).

9. Dang, C. & Jayasena, S. D. Oligonucleotide Inhibitors ofTaqDNA Polymerase Facilitate Detection of Low Copy Number Targets by PCR. J. Mol. Biol. 264, 268–278 (1996).

10. Kubik, M. F., Stephens, A. W., Schneider, D., Marlar, R. A. & Tasset, D. High-affinity RNA ligands to human α-thrombin. Nucleic Acids Res. 22, 2619–2626 (1994).

11. Santana-Viera, L. et al. Combination of protein and cell internalization SELEX identifies a potential RNA therapeutic and delivery platform to treat EphA2-expressing tumors. Mol. Ther. - Nucleic Acids 32, 758–772 (2023).

12. Meyer, S. et al. Development of an Efficient Targeted Cell-SELEX Procedure for DNA Aptamer Reagents. PLOS ONE 8, e71798 (2013).

13. Domenyuk, V. et al. Poly-ligand profiling differentiates trastuzumab-treated breast cancer patients according to their outcomes. Nat. Commun. 9, 1219 (2018).

14. Hornung, T. et al. ADAPT identifies an ESCRT complex composition that discriminates VCaP from LNCaP prostate cancer cell exosomes. Nucleic Acids Res. 48, 4013–4027 (2020).

15. Guido, N., Starostina, E., Leake, D. & Saaem, I. Improved PCR Amplification of Broad Spectrum GC DNA Templates. PloS One 11, e0156478 (2016).

16. Frey, U. H., Bachmann, H. S., Peters, J. & Siffert, W. PCR-amplification of GC-rich regions: ‘slowdown PCR’. Nat. Protoc. 3, 1312–1317 (2008).

17. Sun, D. et al. Computational tools for aptamer identification and optimization. TrAC Trends Anal. Chem. 157, 116767 (2022).

18. Lee, S. J., Cho, J., Lee, B.-H., Hwang, D. & Park, J.-W. Design and Prediction of Aptamers Assisted by In Silico Methods. Biomedicines 11, 356 (2023).

19. Kumar, S. et al. Computational Frontiers in Aptamer-Based Nanomedicine for Precision Therapeutics: A Comprehensive Review. ACS Omega 9, 26838–26862 (2024).

20. Caroli, J., Taccioli, C., De La Fuente, A., Serafini, P. & Bicciato, S. APTANI: a computational tool to select aptamers through sequence-structure motif analysis of HT-SELEX data. Bioinformatics 32, 161–164 (2016).

21. Caroli, J., Forcato, M. & Bicciato, S. APTANI2: update of aptamer selection through sequence-structure analysis. Bioinformatics 36, 2266–2268 (2020).

22. Jiang, P. et al. MPBind: a Meta-motif-based statistical framework and pipeline to Predict Binding potential of SELEX-derived aptamers. Bioinformatics 30, 2665–2667 (2014).

23. Ishida, R. et al. RaptRanker: in silico RNA aptamer selection from HT-SELEX experiment based on local sequence and structure information. Nucleic Acids Res. 48, e82–e82 (2020).

24. Hoinka, J., Zotenko, E., Friedman, A., Sauna, Z. E. & Przytycka, T. M. Identification of sequence– structure RNA binding motifs for SELEX-derived aptamers. Bioinformatics 28, i215–i223 (2012).

25. Asif, M. & Orenstein, Y. DeepSELEX: inferring DNA-binding preferences from HT-SELEX data using multi-class CNNs. Bioinformatics 36, i634–i642 (2020).

26. Zhang, Y., Mo, Q., Xue, L. & Luo, J. Evaluation of deep learning approaches for modeling transcription factor sequence specificity. Genomics 113, 3774–3781 (2021).

27. Song, J. et al. A Sequential Multidimensional Analysis Algorithm for Aptamer Identification based on Structure Analysis and Machine Learning. Anal. Chem. 92, 3307–3314 (2020).

28. Zuker, M. Mfold web server for nucleic acid folding and hybridization prediction. Nucleic Acids Res. 31, 3406–3415 (2003).

29. Biesiada, M., Purzycka, K. J., Szachniuk, M., Blazewicz, J. & Adamiak, R. W. Automated RNA 3D Structure Prediction with RNAComposer. Methods Mol. Biol. Clifton NJ 1490, 199–215 (2016).

30. Zhou, Q., Xia, X., Luo, Z., Liang, H. & Shakhnovich, E. Searching the Sequence Space for Potent Aptamers Using SELEX in Silico. J. Chem. Theory Comput. 11, 5939–5946 (2015).

31. Hiller, M., Pudimat, R., Busch, A. & Backofen, R. Using RNA secondary structures to guide sequence motif finding towards single-stranded regions. Nucleic Acids Res. 34, e117–e117 (2006).

32. Trott, O. & Olson, A. J. AutoDock Vina: improving the speed and accuracy of docking with a new scoring function, efficient optimization, and multithreading. J. Comput. Chem. 31, 455–461 (2010).

33. Abraham, M. J. et al. GROMACS: High performance molecular simulations through multi-level parallelism from laptops to supercomputers. SoftwareX 1–2, 19–25 (2015).

34. Oikonomou, E. D. et al. How natural language processing derived techniques are used on biological data: a systematic review. Netw. Model. Anal. Health Inform. Bioinforma. 13, 23 (2024).

35. Abramson, J. et al. Accurate structure prediction of biomolecular interactions with AlphaFold 3. Nature 630, 493–500 (2024).

36. Rives, A. et al. Biological structure and function emerge from scaling unsupervised learning to 250 million protein sequences. Proc. Natl. Acad. Sci. U. S. A. 118, e2016239118 (2021).

37. Asgari, E. & Mofrad, M. R. K. Continuous Distributed Representation of Biological Sequences for Deep Proteomics and Genomics. PLOS ONE 10, e0141287 (2015).

38. Yang, K. K., Wu, Z., Bedbrook, C. N. & Arnold, F. H. Learned protein embeddings for machine learning. Bioinformatics 34, 2642–2648 (2018).

39. Iwano, N., Adachi, T., Aoki, K., Nakamura, Y. & Hamada, M. Generative aptamer discovery using RaptGen. Nat. Comput. Sci. 2, 378–386 (2022).

40. Wang, Z., et al. AptaDiff: de novo design and optimization of aptamers based on diffusion models. *bioRxiv* 2023.11.25.568693 (2024) doi:10.1101/2023.11.25.568693.

41. Bepler, T. & Berger, B. Learning Protein Sequence Embeddings Using Information from Structure. (2019).

42. Huang, R.-H., Fremont, D. H., Diener, J. L., Schaub, R. G. & Sadler, J. E. A structural explanation for the antithrombotic activity of ARC1172, a DNA aptamer that binds von Willebrand factor domain A1. Struct. Lond. Engl. 1993 17, 1476–1484 (2009).

43. Cooney, K. A. et al. The molecular defect in type IIB von Willebrand disease. Identification of four potential missense mutations within the putative GpIb binding domain. J. Clin. Invest. 87, 1227–1233 (1991).

44. Ruggeri, Z. M. Type IIB von Willebrand disease: a paradox explains how von Willebrand factor works. J. Thromb. Haemost. JTH 2, 2–6 (2004).

45. McInnes, L., Healy, J. & Melville, J. UMAP: Uniform Manifold Approximation and Projection for Dimension Reduction. *ArXiv E-Prints* arXiv:1802.03426 (2018) doi:10.48550/arXiv.1802.03426.

